# Diazo-carboxyl click chemistry enables rapid and sensitive quantification of carboxylic acid metabolites

**DOI:** 10.1101/2023.05.11.540288

**Authors:** Cong Li, Kunlun Cheng, Qijin Zhao, Li Jin, Xuelian Wang, Tongling Liufu, Xutong Zhao, Xiaochuan Li, Xiao Wang, Jia Lyu, Dong Huang, Pingping Li, Xiao-Wei Chen, Zhaoxia Wang, Xinli Hu, Li Quan, Zhixing Chen

## Abstract

Carboxylic acids are central metabolites in bioenergetics, signal transduction and post-translation protein regulation. Unlike its genomic and transcriptomic counterparts, the quest for metabolomic profiling in trace amounts of biomedical samples is prohibitively challenging largely due to the lack of sensitive and robust quantification schemes for carboxylic acids. Based on diazo-carboxyl click chemistry, here we demonstrate DQmB-HA method as a rapid derivatization strategy for the sensitive analysis of hydrophilic, low-molecular-weight carboxylic acids. To the investigated metabolites, DQmB-HA derivatization method renders 5 to 2,000-fold higher response on mass spectrometry along with improved chromatographic separation on commercial UHPLC-MS machines. Using this method, we present the near-single-cell analysis of carboxylic acid metabolites in mouse egg cells before and after fertilization. Malate, fumarate and β-hydroxybutyrate were found to decrease in mouse zygotes. We also showcase the kinetic profiling of TCA-cycle intermediates inside adherent cells cultured in one well of 96-well plates during drug treatment. FCCP and AZD3965 were shown to have overlapped but different effects on the isotope labeling of carboxylic acids. Finally, we apply DQmB-HA method to plasma or serum samples (down to 5 μL) from mice and humans collected on pathological and physiological conditions. The measured changes of succinate, β-hydroxybutyrate, and lactate in blood corroborate previous literatures in ischemia-reperfusion injury mouse model, acute fasting-refeeding mouse model, and human individuals diagnosed with mitochondrial dysfunction diseases, respectively. Overall, DQmB-HA method offers a sensitive, rapid and user-friendly quantification scheme for carboxylic acid metabolites, paving the road toward the ultimate goals of single-cell metabolomic analysis and bedside monitoring of biofluid samples.

## Introduction

Carboxylic acid metabolites are mainstays in core metabolic pathways including the tricarboxylic acid (TCA) cycle, redox homeostasis^1^, fatty acid metabolism, and amino acid metabolism^2,3^. They compose the dynamic metabolome that fuels essentially all the biochemical processes. Studies of individual carboxylic acid metabolites have recently unveiled many new roles these metabolites play in both known and unknown biochemical pathways^4–12^. At the same time, monitoring the dynamics of carboxylic acids as part of a comprehensive metabolites pool provides a quantitative, holistic understanding of cellular metabolism^13–20^. Recently the rapid growth of single-cell technologies is promising the dawn of single-cell omics, where (epi)genomic, transcriptomic, proteomic, and metabolomic data can be integrated and analyzed to provide an unprecedented level of information^21–26^. This picture, however, is bottlenecked at the metabolomic analysis which, unlike the amplifiable nucleic acids that can be thoroughly sequenced, still suffers a low signal sensitivity. Cutting-edge mass spectrometry technologies have been carried out for single-cell or even single-organelle analysis of selected groups of biomolecules^27–38^, but the state-of-the-art metabolomic analysis of carboxylates mostly hinges on hydrophilic interaction liquid chromatography (HILIC) or reverse phase liquid chromatography (RPLC) coupled with mass spectrometry (MS) technologies which give abundant chemical information with molecular specificity. Particularly, carboxylic acid metabolites with less than six carbon atoms are of low molecular weights, extremely hydrophilic, and negatively charged, which altogether prompts challenges regarding separation in liquid chromatography and ionization detection in mass spectrometry. Consequently, the LC-MS analysis of carboxylic acid metabolites is routinely carried out using samples containing 10-100 pmol metabolites, which is far away from high-throughput micrometric analysis, let alone single cell-level detections.

A promising strategy to increase the chemical stability and detection sensitivity of carboxylic acids is chemical derivatization, which masks the cumbersome carboxyl group and adds auxiliary functional groups to aid chromatographic separation and to switch negative ionization to positive ionization which is more sensitive in MS. Among common carboxyl-reactive reagents, amines^39–41^, hydrazines^42–44^, hydroxylamine^45^, and alcohols^46^ are the most popular chemicals used to derivatize carboxylic acids via condensation reactions^47^. However, these coupling reactions typically require excess coupling reagents to drive derivatization. During derivatization, coupling reagents inevitably release alkylated urea or oxophosphorous(V) byproducts^48^, which are often removed by additional extraction and concentration steps, complicating the overall workflow^41,45^. Reactive bromides and triflates have also been suggested as derivatization reagents^49^. These electrophiles generally exhibit low reactivity and often require excess bases or heating up to 90 °C to facilitate derivatization^50^, causing degradation of multiple metabolites such as β-ketoacids^51^. Diazo compounds have long been known to react with carboxylic acids and serve as valuable esterification reagents in organic synthesis^52^. Although fluorescent diazomethyl compounds were already applied to the derivatization of fatty acids back in the 1980s^53–55^, diazo compounds have not been impacting biomedical research until a recent resurgence^56^. With the successful demonstration of diazo-based click bioconjugation in protein delivery^57,58^, nucleic acid modification^59,60^, and protein crosslinking^61,62^, we envision this chemistry is particularly well-suited in derivatizing carboxylic acid metabolites due to its selective reactivity and mild procedure.

Here, we present a derivatization strategy for versatile and sensitive metabolomics analysis of general hydrophilic carboxylic acids in biological samples using LC-MS/MS. This method is based on a newly designed, stable, carboxyl-reactive diazo reagent quinolinyl diazo(*p*-methyl)phenylacetate (DQmB) and an orthogonal, ketone-reactive hydroxylamine (HA). Using a combined derivatization kit, carboxylic acids in aqueous samples are handily converted to DQmB esters with enhanced chromatographic separations and boosted signal on mass spectrometers, giving up to three orders of magnitude higher sensitivity. We showcase five applications on bioanalysis, including near-single-cell metabolic profiling in mouse zygotes, metabolic flux analysis using neonatal rat primary cardiomyocytes cultured in 96 well-plate, serum analysis in ischemia-reperfusion and fasting-refeeding mouse models, and comparative study of clinical samples between mitochondrial myopathy patients and healthy volunteers.

## Results

### Design and synthesis of DQmB

Diazo compounds react with carboxylic acids under acidic to neutral conditions in a traceless fashion, in which no additives are needed and no other byproducts than N_2_ are released. Encouraged by the unique properties of diazo-carboxylic acid chemistry, we chose diazo phenylacetate ester as our scaffold due to its synthetic convenience enabled by well-established diazo transfer reactions. We appended an additional methyl group at the para position of phenylacetate based on two reasons: a) *p*-methyl group increases the reactivity towards carboxylic acids and at the same time keeps a low hydrolysis/esterification ratio^63^. b) phenyl group is ubiquitous in all biological milieu, which makes it an undesirable target ion in the tandem mass spectrum. In the contrast, the *p*-methylphenyl group is sparely found in biological and clinical samples. The selection of *p*-methylbenzylic carbocations as product ions would specifically pick out all the derivatized TCA intermediates and thus reduce false positive mass signals. To further improve mass response in positive mode, we reasoned that the addition of a nitrogen-containing group, such as the quinolinyl group, could promote protonation and increase ionization efficiency. Finally, quinolinylmethyl diazo(*p*-methyl)phenylacetate (DQmB) was presented for the derivatization of carboxylic acids (Fig. 1a). DQmB was readily synthesized starting from two-step esterification of *p*-methylphenyl acetic acid, followed by one-step diazo transfer with an overall yield of 46% in three steps (See SI Chemical synthesis of DQmB).

**Figure 1.**
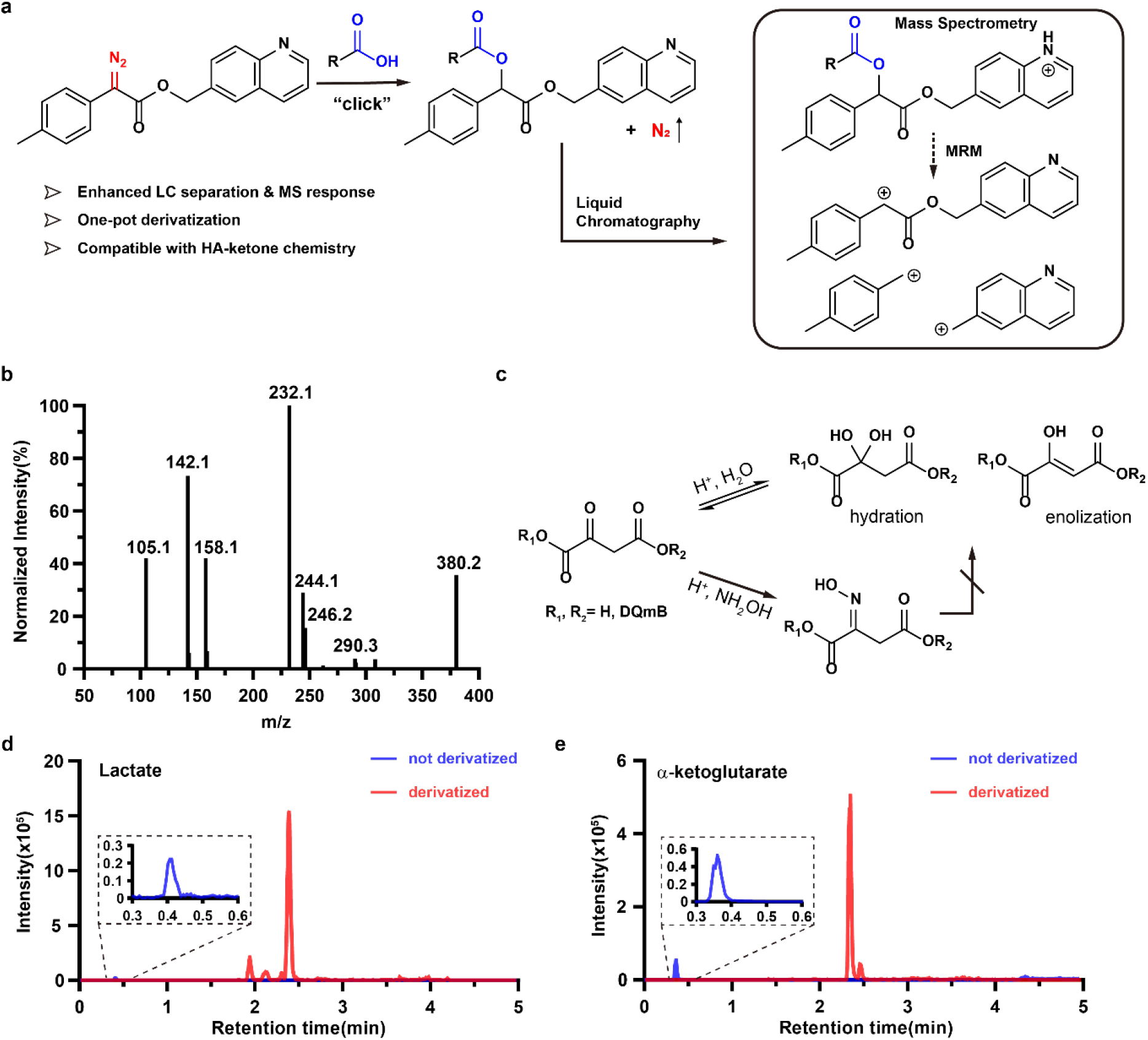
DQmB-HA derivatization method, based on diazo-click chemistry, enables sensitive quantification of carboxylic acid metabolites using LC-MS/MS. a, Schematic of DQmB derivatization chemistry and the sensitive mass spectrometry reactions of analytes. MRM, multiple reaction monitoring. b, Secondary mass spectrum of lactate after DQmB derivatization. c, Hydroxylamine (HA) reacts with α-keto acids or esters to form stable oximes, inhibiting hydration or enolization pathways of α-keto acids or esters. d, e, Overlayed chromatograms of Lactate(d) and α-ketoglutarate(e) with and without derivatization. Lactate was derivatized with only DQmB, and α-ketoglutarate was derivatized with both DQmB and HA. Not derivatized: [Lactate]=100 μM, [α-ketoglutarate]=10 μM, negative mode. Derivatized: [Lactate]=10 μM, [α-ketoglutarate]=10 μM, positive mode.

### Development of DQmB-HA derivatization method

With DQmB in hand, we set out to aquire mass spectrometric parameters for carboxylic acid metabolites (Table S1) and found that carboxylic acids were readily derivatized by DQmB in a mixture of water and acetonitrile. As a typical example, the secondary mass spectrum of L-lactate after DQmB derivatization showed the formation of the derivatized product Lac-DQmB, which had a molecular weight of 379.2 Da (Fig. 1b). The peaks at 142.1 and 105.1 corresponded to quinolinylmethyl and benzyl cations respectively, both of which belonged to fragment ions from Lac-DQmB as we expected (Fig. S1). Next, we optimized the derivatization conditions for carboxylic acids by investigating the effects of temperature, concentration, and water content on derivatization. At temperatures below 55 °C, carboxylic acids were barely derivatized while the extent of derivatization increased rapidly at temperatures above 55 °C, and reached a maximum at 70 °C (Fig. S2a). In contrast, DQmB concentration had a limited, if any, effect on derivatization (Fig. S2b). Since acetonitrile was selected as a co-solvent to increase the solubility of DQmB, we next screened a series of water-to-acetonitrile ratios. To our delight, DQmB derivatization was highly tolerant to water (Fig. S3), making it intrinsically a suitable derivatization reagent for analyzing biosamples. In addition, aqueous hydrochloride acid (HCl) was added to ensure the protonation of carboxylic acids for the reaction with DQmB. A broad range of HCl concentrations (1-50 mM) was comparably suitable for DQmB derivatization (Fig. S4).

However, under the optimized conditions in the derivatization of L-lactate, α-ketoacids behaved poorly during routine chromatographic separation, possibly due to hydration or enolization of the carbonyl group at α position of the electron-withdrawing ester group (Fig. 1c). To minimize these side reactions, we exploited hydroxylamine (HA) as a carbonyl-reactive group to yield stable oximes with lower hydration or enolization tendency. Finally, by combining DQmB and HA, we tailored a DQmB-HA method to derivatize carboxylic acid metabolites in biological samples. In this updated protocol, 4 volumes of acetonitrile solution containing 20 mM DQmB and 1 mM HA were directly added to 1 volume of aqueous biological samples. Then, 10 volumes of 10 mM aqueous hydrochloride acid (HCl) were added and the derivatization mixture was incubated at 70 °C for 20 min. Under these conditions, carbonyls were readily converted to oximes, and carboxylates were esterified by DQmB, giving stable adducts with few byproducts for direct LC-MS/MS analysis. In addition, DQmB-HA derivatization method effectively decreased the polarity of carboxylic acids, giving rise to improved retention on reverse phase column and symmetrical chromatography peaks for unambiguous assignments of analytes.

Overall, the improvements of LC behavior and MS response lead to a much higher sensitivity of polar carboxylic acids. Compared with non-derivatized lactate, lactate derivatized by DQmB-HA method had a longer retention time and the intensity of MS response was 750-fold higher (Fig. 1d). The elution of derivatized α-ketoglutarate was similarly delayed compared to non-derivatized α-ketoglutarate and the intensity of MS response was enhanced by approximately 10 folds (Fig. 1e). Generally, DQmB-HA method enhanced MS signal intensity of carboxylic acid metabolites by 5-2000 folds and reduced the lower limit of detection (LOD) for investigated carboxylic acids to the level of 1-10 femtomoles, which is comparable to oBHA method^45^ in our hands (Table S2). However, oBHA method took more than 80 min to complete the whole derivatization process. In contrast, DQmB-HA method drastically simplified the procedures of sample preparation and the overall derivatization process took less than 30 minutes, which would minimize the potential degradation and the adsorption onto lab plasticwares of carboxylic acids. Therefore, it is possible to analyze carboxylic acid metabolites with DQmB-HA method in trace amounts of biomedical samples.

### Analysis of carboxylic acid metabolites in scant cells

Cellular transcriptomics, epigenomics, and proteomics undergo remarkable changes during the early stage of mammalian embryogenesis. However, our understandings regarding the dynamic changes of metabolites from oocytes to zygotes remain limited due to the scarcity of samples. To test the power of our method, scant numbers of mouse oocytes or zygotes were handpicked into 200 μL tubes using micro capillaries and immediately derivatized by DQmB-HA (Fig. 2a). The derivatized samples were directly injected into LC-MS/MS system without further processing, minimizing the loss or the dilution of the metabolites.

**Figure 2.**
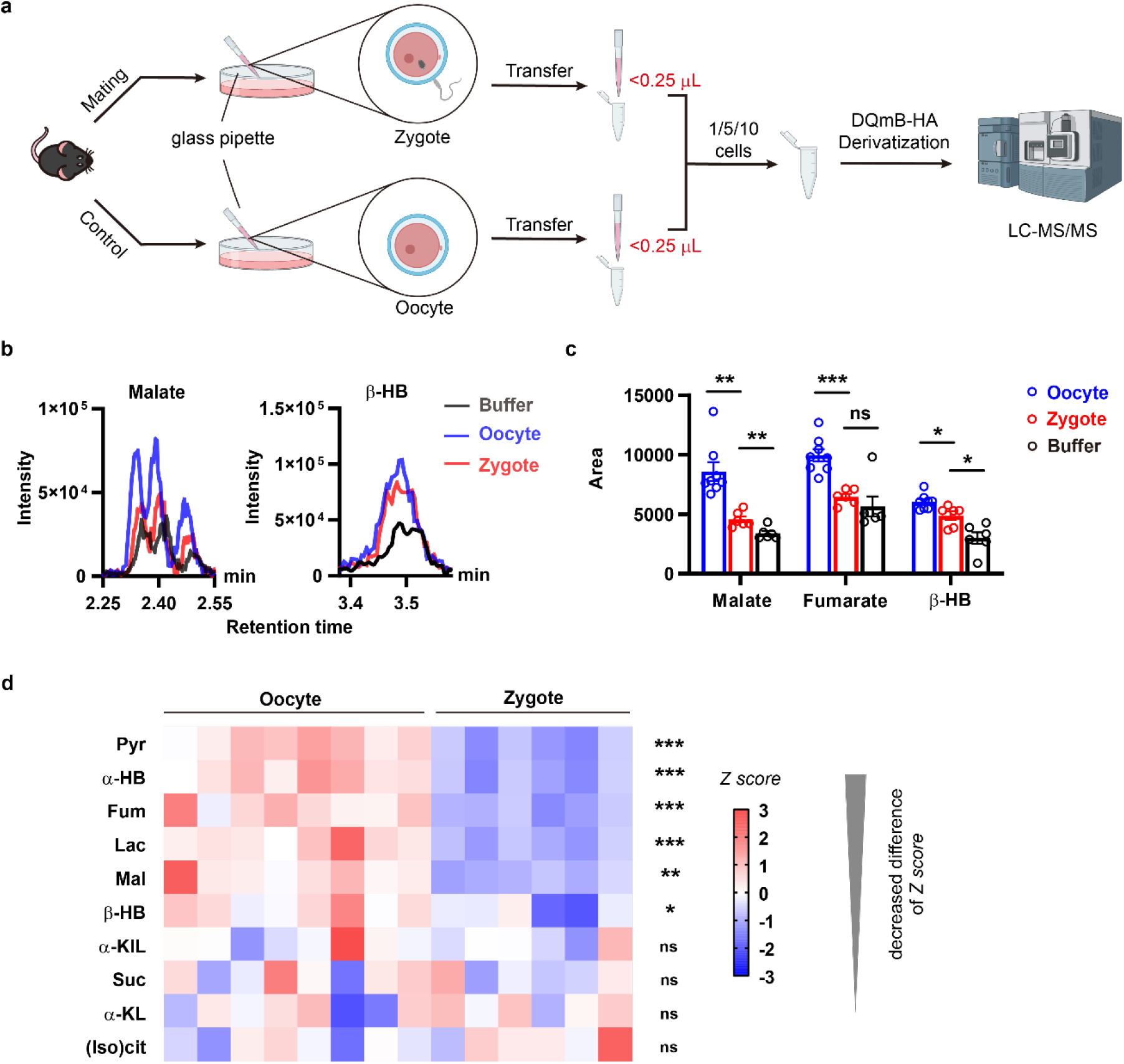
Fertilization remodeled levels of carboxylic acid metabolites in mouse zygotes as revealed by DQmB-HA derivatization of scant cells. a, Workflow for the quantitative profiling of carboxylic acid metabolites in mouse oocytes and zygotes using DQmB-HA derivatization method. b, Representative chromatograms of malate and β-hydroxybutyrate in buffer, 10 oocytes or 10 zygotes. c, Comparisons of malate, fumarate, β-hydroxybutyrate in 10 oocytes, 10 zygotes, and buffer. Data are mean ± s.e.m.; unpaired Student’s *t* test (n=8 for Oocyte group, n=6 for Zygote group, n=6 for Buffer group). d, Heat map showing Z scores of carboxylic acid metabolites in 10 zygotes (n=6) or 10 oocytes (n=8). The rows are arranged according to the differences of Z scores. Unpaired Student’s *t* test (n=8 for Oocyte group, n=6 for Zygote group).

In order to determine the minimal number of cells needed as input, we titrated the amounts of carboxylic acids in 1, 5, and 10 cells using DQmB-HA method. For both oocytes and zygotes, the MS signals increased as the number of cells increased (Fig. S7a-d). Compared to 1-cell samples, 10-cell samples and 5-cell samples had similar numbers of significant events detected in total (10 for 10-cell, 9 for 5-cell) (Fig. S7e, g). Specifically, for 10-cell samples, half of the detected significant events were in Oocyte group and another half were in Zygote group. In contrast, for 5-cell samples, most of the detected significant events were in Oocyte group and only one was in Zygote group, indicating an uneven distribution of statistical power between Oocyte group and Zygote group. According to the signs of differences, we further categorized the detected significant events into increased or decreased. For 10-cell samples, only 2 out of 10 significant events were decreased events, while for 5-cell samples, 4 out of 9 significant events were decreased events (Fig. S7f, h). Therefore, samples with 10 cells were selected as input in the following experiments to ensure sufficient statistical power and reduced false negative results.

The characteristic peaks belonging to each carboxylic acid were identified in the chromatograms of 10-cell samples (Fig. 2b, Fig. S8a). For certain carboxylic acids, peaks could be directly compared to visualize the differences between 10 zygotes and buffer, which was consistent with statistical results (Fig. S8b-d). Quantitatively, malate, fumarate and β-hydroxybutyrate decreased by about 80%, 80% and 40%, respectively (Fig. 2c), as validated in at least two out of three independent experiments (Table S3). On the other hand, the levels of succinate, α-ketoleucine and (iso)citrate did not change significantly in mouse zygotes (Fig. 2d, Table. S3). Our results indicated that the drastic changes of carboxylic acids can be detected using the DQmB-HA derivatization in samples containing down to 10 egg cells. The fertilization remodeled levels of carboxylic acid metabolites in mouse zygotes, providing a metabolomic insight on the early embryogenesis.

### Isotope tracing of cellular carboxylic acid metabolites in 96-well plates

Isotope tracing is a powerful tool to provide quantitative information about metabolic dynamics in cells under essentially no chemical perturbation. To obtain robust results in isotope tracing experiments, however, one has to prepare samples with a larger number of cells than those in non-tracing experiments due to the isotope dilution during labeling. This seems especially challenging when cell numbers are limited, i.e., in certain primary cell lines or under a specific cell phase. In addition, high-throughput isotope tracing experiments at multiple time points require additional groups of cells, further impeding the large-scale implementation of isotope tracing experiments in applications such as drug screening. Featured by high sensitivity and concise workflow, DQmB-HA derivatization is promising in minimizing both cell and reagent consumptions for advanced isotope tracing experiments.

To demonstrate the feasibility of isotope tracing experiments in one well of 96-well plates, U-^13^C_6_-glucose was employed as the sole C-13 source to initiate isotope tracing in neonatal rat ventricular myocytes (NRVMs).

In a modified DQmB-HA derivatization protocol, DQmB and HA were directly added to adherent NRVMs cultured in wells, followed by ultrasonic detachment/lysis of cells and further addition of aqueous hydrochloride acid (Fig. 3a). In control NRVMs, there was a time-dependent global accumulation of ^13^C-labeled carboxylic acid metabolites. Among them, ^13^C-labeled pyruvate and lactate accumulated most rapidly, reaching ∼60% and ∼40% labeled fractions within 10 minutes, respectively (Fig. 3b, Fig. S9a). At the entry of TCA cycle, (iso)citrates were ^13^C-labeled at a slightly slower rate of ∼30% fraction in 10 minutes (Fig. 3b). Other downstream carboxylic acids, such as α-ketoglutarate, were ^13^C-labeled at even slower rates (Fig. S9b-d). In parallel, NRVMs were treated with classic respiration inhibitors, and their effects on ^13^C-labeling kinetics during a 60-minute time course were traced. The treatment with oligomycin or combined rotenone/antimycin A deaccelerated ^13^C-labeling rates of (iso)citrate and fumarate in NRVMs, compared with control NRVMs (Fig. 3c, Fig. S10a, b). In contrast, the treatment with FCCP increased ^13^C-labeled fractions of α-ketoglutarate, succinate, and fumarate by 10%-20% of the absolute total amount (Fig. 3c, Fig. S10c-e). FCCP boosted the TCA flux so much that (iso)citrate were even ^13^C-labeled to ∼20% in a second run as the M+4 fraction was observed to increase continuously and the M+2 fraction peaked at 5 min before starting to decrease (Fig. 3c, Fig. S10f, g).

**Figure 3.**
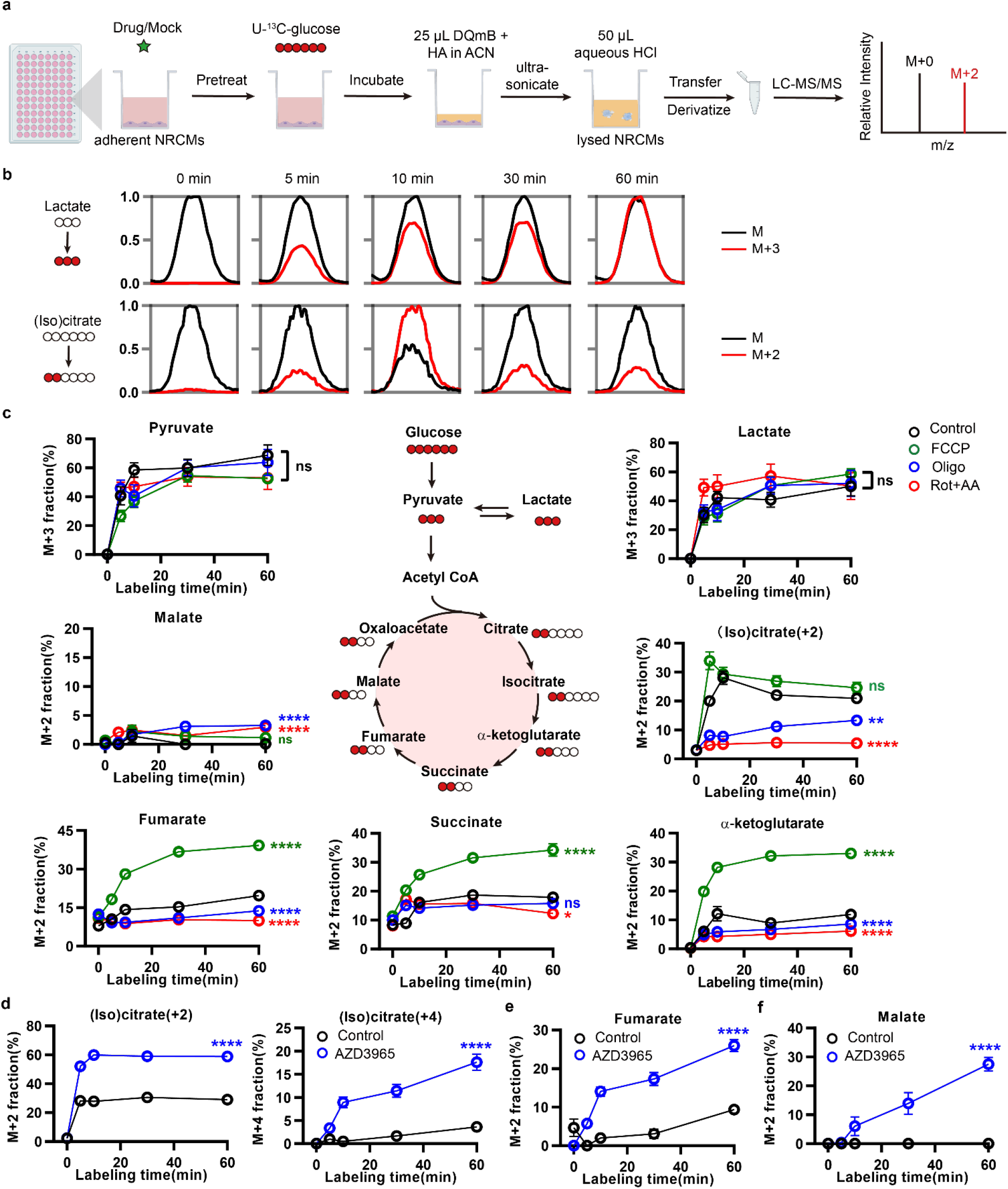
Kinetic isotope tracing in 96-well plates reveals distinct effects of small molecules on ^13^C-labeling rates of carboxylic acid metabolites by U-^13^C-glucose in neonatal rat ventricular myocytes (NRVMs). a, Workflow for the kinetic isotope tracing experiments using U-^13^C_6_-glucose in 96-well plate formats integrated with DQmB-HA derivatization method. b, Representative merged chromatograms showing both labeled and unlabeled fractions of lactate and (iso)citrate at multiple time points in control NRVMs. c, Comparisons of the labeled fractions of carboxylic acids at 0, 5, 10, 30, and 60 minutes in NRVMs treated with FCCP, Oligo, or Rot+AA separately, to control. FCCP: carbonyl cyanide-*p*-trifluoromethoxyphenylhydrazone, Oligo: Oligomycin, Rot: Rotenone, AA: Antimycin A. Data are mean ± s.e.m.; one-way ANOVA followed by multiple comparisons (n=8 for each group). d-f, Comparisons of the labeled fractions of (iso)citrate(d), fumarate(e), and malate(f) at 0, 5, 10, 30, and 60 minutes in NRVMs treated with AZD3965 to control. Data are mean ± s.e.m.; unpaired Student’s *t* test (n=8 for each group).

Next, we showcased that DQmB-HA method could facilitate the exploration of possible biochemical perturbations induced by small molecules to the central carbon metabolism. AZD3965 is a selective inhibitor for monocarboxylate transporter 1(MCT-1) as well as an anti-cancer drug candidate in early-phase clinical trials^62–64^(Fig. S11a). We used AZD3965 to treat NRVMs to evaluate the effect of MCT-1 on the metabolic flow in TCA cycle. Similar to FCCP treatment, AZD3965 treatment significantly accelerated ^13^C-labeling kinetics of (iso)citrate, fumarate, and malate (Fig. 3d-f), while showing nearly no effects on lactate and pyruvate (Fig. S11b, c). However, AZD3965 showed little effect on ^13^C-labeling of succinate within 60 minutes as AZD3965 works in a different mechanism compared to FCCP (Fig. S11d, e). Overall, these data support that the DQmB-HA derivatization method offers a much higher throughput in kinetic isotope tracing experiments, scaling down the previously costly assay to 96-well plates (∼1×10^5^ cells)..

### Monitoring carboxylic acid metabolites in peripheral blood of two mouse models

DQmB-HA derivatization method is not only suitable for cellular metabolomic analysis but also generally applicable to body fluids. We showcased two examples of quantitatively analyzing carboxylic acid metabolites in blood. Succinate is a recently reported biomarker of myocardial ischemia-reperfusion (IR) injury, exhibiting over a 3-fold increase in peripheral regions at risk^4^. With the DQmB-HA derivatization method, we aim to monitor the dynamics of carboxylic acid metabolites, especially succinate, in peripheral blood during the entire process of myocardial IR injury. At each time point, 50 μL of blood was collected from mouse tail tips and 5 μL plasma was subjected to DQmB-HA derivatization (Fig. 4a). Our results showed that succinate was accumulated in plasma at 30 minutes after ischemia in both Ischemic&Reperfused (IR) group and Ischemic (I) group, and dropped back to basal levels at 24 hours after reperfusion in IR group (Fig. 4b, c, Fig. S12a). Though succinate was also slightly accumulated in Sham group, the increase was smaller than those in IR group and I group (Fig. 4c, Fig. S12a, b). Among other investigated metabolites, the levels of α-hydroxybutyrate and (iso)citrate in plasma remained nearly unchanged during the first 5 minutes after reperfusion but decreased to basal levels at 24 hours after reperfusion (Fig. 4d-f). Therefore, DQmB-HA derivatization method renders an appealing opportunity to fully profile carboxylic acid metabolites during the whole process of IR injury.

**Figure 4.**
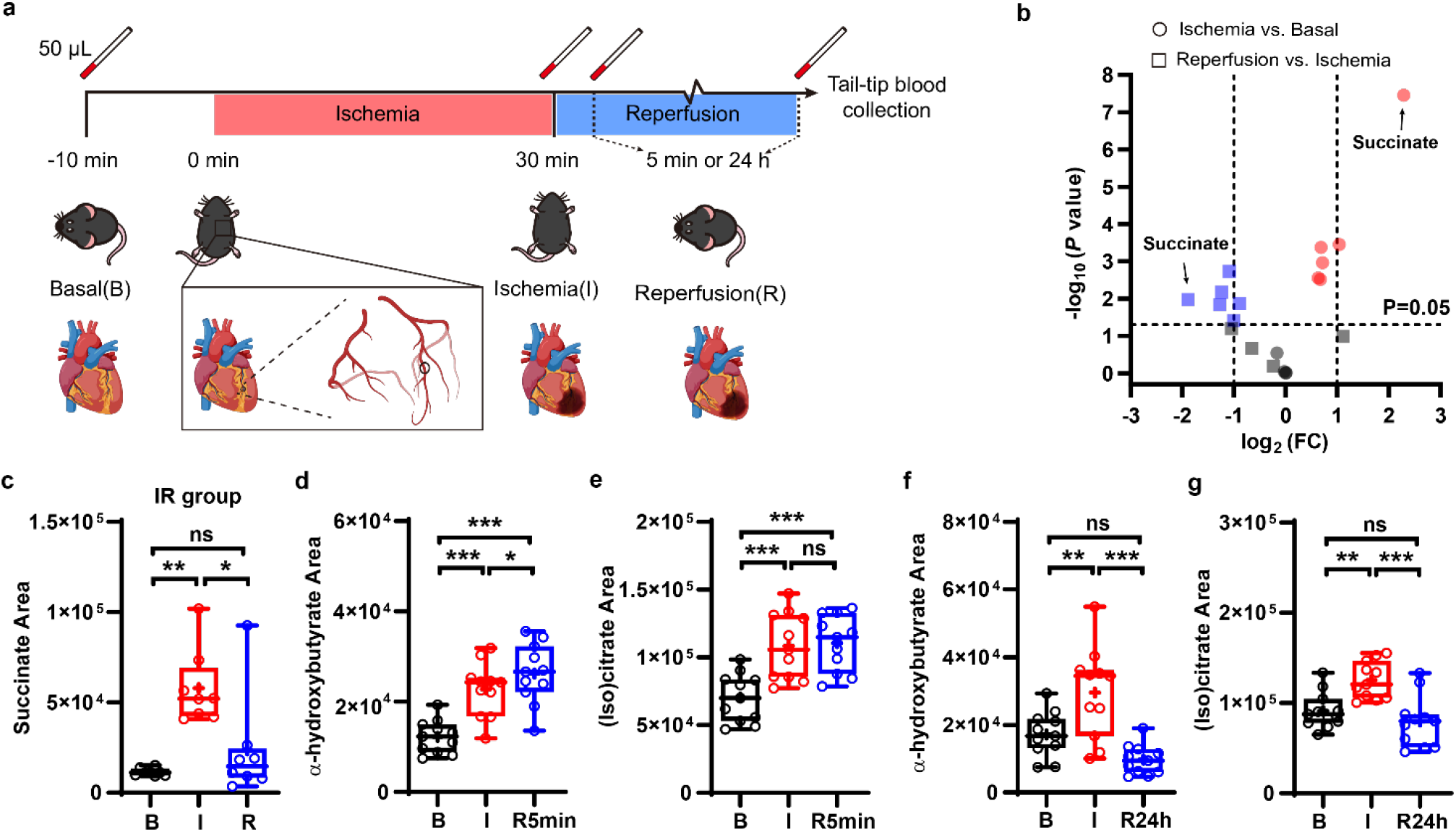
The minimal consumption of serum sample (5 μL) used in DQmB-HA analysis enables temporal monitoring of carboxylic acid metabolites during myocardial ischemia-reperfusion injury. a, Schematic of monitoring the levels of carboxylic acid metabolites in mouse serum before myocardial ischemia (Basal), at 30 minutes post myocardial ischemia (Ischemia) and at 5 min or 24 hours post reperfusion (Reperfusion). b, Volcano plot showing correlations between P values (evaluated by two-tailed ratio paired *t* test, n=8) and relative areas (Fold Changes, FC) for all investigated carboxylic acids at Ischemia (relative to at Basal) and at Reperfusion (relative to at Ischemia) shown in (a) in IR group. c, Comparisons of the measured levels of succinate in mouse serum at Basal (B), Ischemia (I), and Reperfusion (R) in IR group. Data are mean ± s.e.m.; repeated measures one-way ANOVA followed by multiple comparisons (assume sphericity, n=8 for each group). d-g, Dynamics of α-hydroxybutyrate and (iso)citrate in mouse serum during myocardial ischemia-reperfusion injury. R5min represents 5 min post reperfusion in d, e and R24h represents 24 h post reperfusion in f, g. Data are mean ± s.e.m.; repeated measures one-way ANOVA followed by multiple comparisons (assume sphericity, n=11 for each group).

In another model, we aim to investigate the effects of chronic diets on ATP-generating metabolism in mice during the acute fasting-refeeding process. DQmB-HA derivatization was employed to monitor the levels of carboxylic acid metabolites in plasma collected from mice chronically fed with either a normal chow diet (NC) or a high-fat diet (HFD). Corroborating the literature^64,65^, acute fasting induced a drastic increase of β-hydroxybutyrate in both NC mice and HFD mice, while refeeding restored the levels of β-hydroxybutyrate (Fig. 5b, c). At all three investigated time points, the levels of β-hydroxybutyrate in HFD mice were more discrete than those in NC mice, suggesting that the chronic diets might contribute to metabolic heterogeneity. Other ketone body metabolites such as α-hydroxybutyrate, α-ketoleucine, α-ketoisovalerate, and α-ketoisoleucine, also increased after acute fasting and decreased after refeeding in NC mice (Fig. S13a-d). Yet no significant differences of α-ketoleucine, α-ketoisovalerate, and α-ketoisoleucine during the whole process were observed in HFD mice (Fig. S13e-h). In contrast, the levels of lactate in NC mice first decreased after 12-h acute fasting and then increased to higher levels than those in basal after 6-h refeeding (Fig. 5d), while in HFD mice the levels of lactate were not affected (Fig. 5e). Similarly, the levels of pyruvate kept nearly unchanged after acute fasting but accumulated to reach much higher levels than those in basal (Fig. S13i, j). These results indicated that the chronic diet condition has a profound impact on the metabolism of carboxylic acids during the acute fasting-refeeding process. The minimal sample requirement of DQmB-HA derivatization method allowed the continuous monitoring of individual animals during the acute fasting-refeeding process, paving the way to perform additional investigations with precious experimental subjects, such as genetically modified animals.

**Figure 5.**
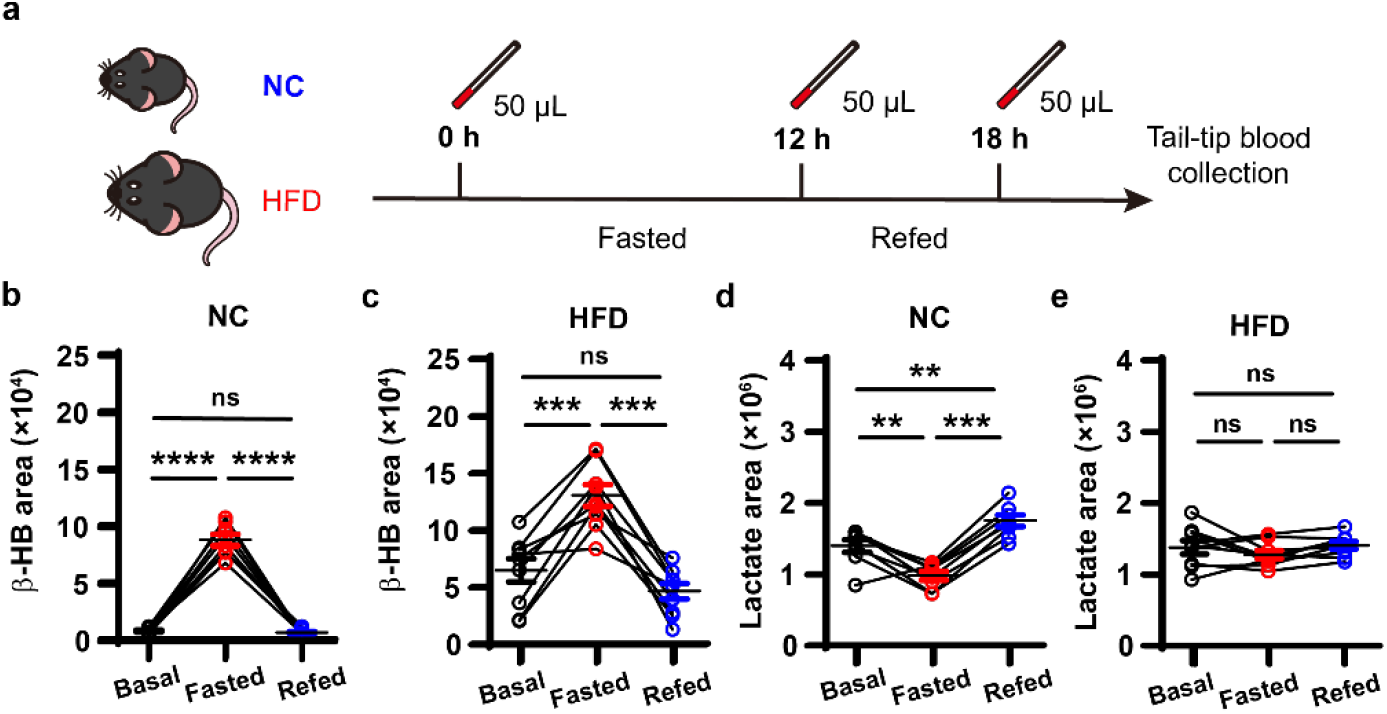
Chronic diet styles affect the dynamic changes of carboxylic acid metabolites in mouse serum during the acute fasting-refeeding process. a, Workflow for monitoring carboxylic acid levels in mouse serum during the fasting-refeeding process. b-e, Dynamic changes of β-hydroxybutyrate (b, c) and lactate (d, e) in serum from normal chow-diet mouse (NC) or high-fat-diet mouse (HFD) before fasting (Basal), after 12-h fasting (Fasted) and after 6-h refeeding (Refed). Data are mean ± s.e.m.; repeated measures one-way ANOVA followed by multiple comparisons (assume sphericity, n=8 for NC group, n=9 for HFD group).

### Profiling metabolic features of carboxylic acid metabolites in clinical samples

Encouraged by results from blood samples in two mouse models, we sought to apply DQmB-HA derivatization method to human blood samples in a clinical setting. Mitochondrial diseases are a large group of metabolic disorders with various genetic causes, all of which in the end lead to mitochondrial dysfunction. We focused on a specific type of mitochondrial disease: mitochondrial encephalomyopathy, lactic acidosis, and stroke-like episodes (MELAS) syndrome. The clinical samples were collected by intravenous blood collection, after which 5 μL plasma or serum was derivatized by DQmB-HA method and then subjected to further analysis (Fig. 6a).

**Figure 6.**
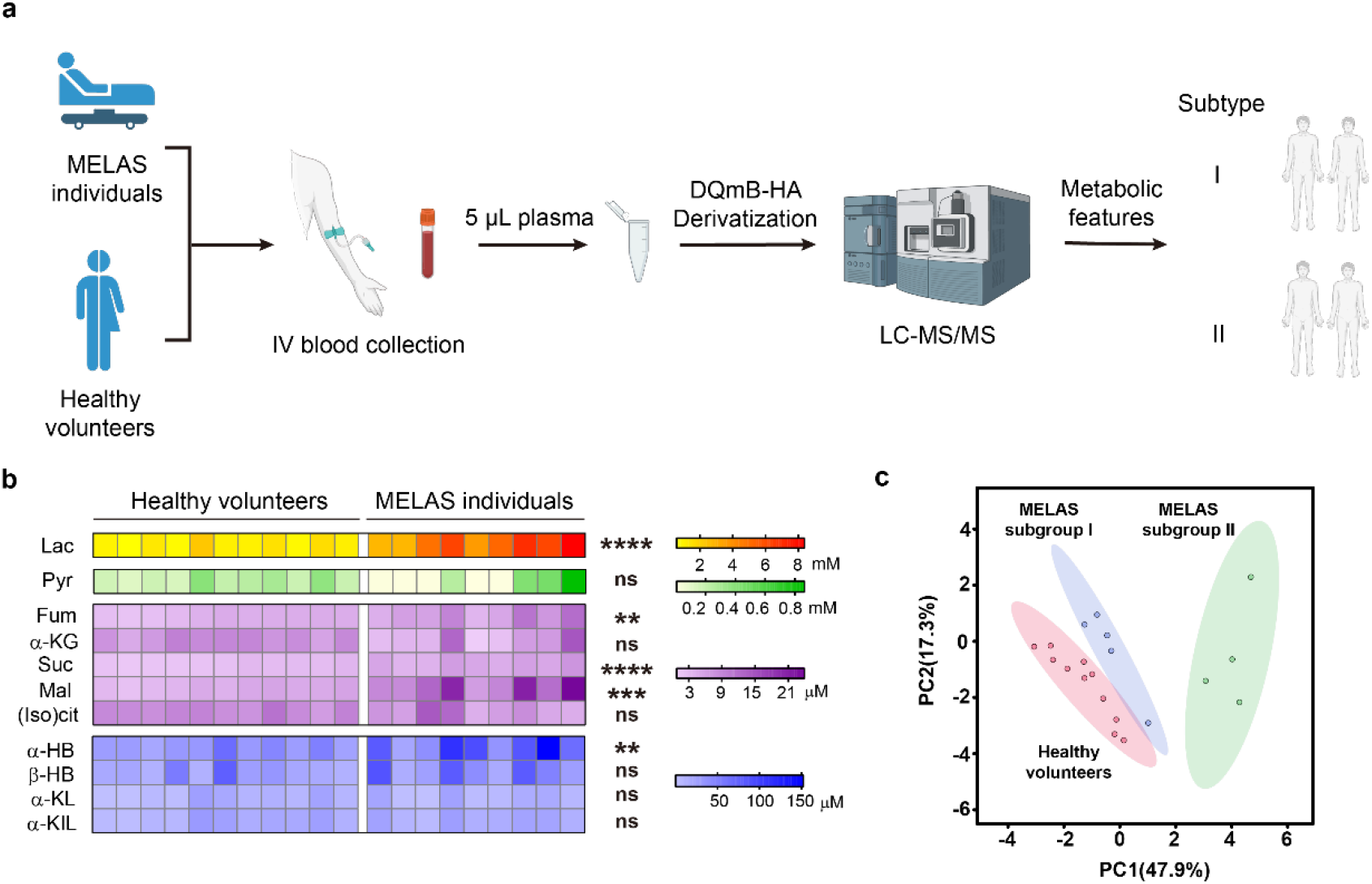
DQmB-HA method enables profiling carboxylic acid metabolites in clinical blood samples. a, Schematic of simple and rapid monitoring of carboxylic acid levels in blood samples of individuals diagnosed with mitochondrial diseases and healthy volunteers. b, Heat map showing concentrations of carboxylic acid metabolites in blood samples from healthy volunteers (11 plasma samples) and MELAS individuals (6 serum samples, 3 plasma samples). Unpaired Student’s *t* test (n=11 for Healthy volunteers group and n=9 for MELAS individuals group). c, Two-dimension (2D) scores plot of carboxylic acid metabolites in blood samples from healthy volunteers (n=11) and MELAS individuals (n=9). Ellipses represent 95% confidence regions.

11 healthy volunteers and 9 MELAS individuals were enrolled in this study (Table S4, S5).Compared to healthy volunteers, MELAS individuals had higher lactate concentrations in their blood calculated from standard calibration curves (Fig. 6b, Fig S14a), which was consistent with previous reports^66^. We also noticed the moderate accumulation of fumarate, succinate, malate, and α-hydroxybutyrate in blood samples from MELAS individuals (Fig. 6b, Fig. S14b-k). The principle component analysis (PCA) plot showed that MELAS individuals involved in this study could be further divided into two subgroups designated randomly as I and II (Fig. 6c). Principle component 1(PC1) accounted for major differences (47.9%) between subgroup I and subgroup II. The loadings of PC1 indicated no single metabolite as dominant, but instead, showed comparable contributions from fumarate, malate, succinate, lactate, and α-hydroxybutyrate (Table. S6), which manifested the high metabolic complexity of MELAS syndrome. Interestingly, along the PC1 axis, MELAS subgroup I was closer to healthy volunteers, suggesting that subgroup I should have some similar metabolic features as healthy volunteers have, which may be a hint about providing precision treatment for MELAS individuals.

Above results suggested that, although MELAS individuals were clinically diagnosed with the same syndrome and even sequenced to have the same genetic mutations (Table S4), their metabolic features could be different and further subtyped using metabolomics data enabled by microanalysis, which may benefit future clinical research for MELAS syndrome.

## Discussion

Inspired by previous work on diazo compounds as carboxyl-derivatization reagents^53–55^, we present DQmB-HA derivatization method for convenient and sensitive analysis of cellular samples and blood samples. The design of DQmB balances the stability and the reactivity of the diazo group, allowing safe and easy handling of derivatization reagents in labs specialized in metabolomics. DQmB enhances both LC separation and MS ionization, yet is synthetically simple and accessible. The bioorthogonality of diazo/carboxyl conjugation ensures the high compatibility of DQmB with other derivatization chemistry such as hydroxylamine-ketone condensation, enabling multiplexed bioanalysis. As DQmB and HA synergize to derivatize carboxyl and ketone groups, DQmB-HA method is devised for the detection of ketoacids. In this study, pyruvate is quantified in generally all types of tested samples, while α-ketoglutarate, α-ketoleucine, α-ketoisoleucine, and α-ketoisovalerate can be well quantified in fresh blood samples and samples containing down to 10^4^ cells. Oxaloacetate, a low-abundance β-ketoacid prone to decarboxylation^67,68^, is not yet detectable in a reliable manner in biological samples using DQmB-HA protocol. Hydrophobic carboxylic metabolites, such as fatty acids, are promising targets for DQmB. The protocols for detecting these hydrophobic carboxylic acids need to be separately designed and would be reported in due course.

DQmB-HA derivatization method is generally applicable to various types of biomedical samples. It particularly benefits experiments that have been previously prohibited by limited amounts of samples and/or complicated sample treatment (e.g. single-cell analysis and clinical sample analysis). For example, single-cell metabolomics generally lagged behind other single-cell omics, largely due to the diverse physiochemical properties of metabolites and the lack of general amplification of small molecules. We demonstrate the dramatic decrease of malate, fumarate, and β-hydroxybutyrate in oocytes after fertilization at the 10-cell level. Therefore, DQmB-HA derivatization method is an important step toward the goal of single-cell carboxylic-acid metabolic analysis. To achieve single-cell carboxylic acid metabolomics, the diazo-carboxyl click chemistry, could serve as a prototype among all kinds of derivatization strategies. In parallel with single-cell analysis, clinical analysis is technically demanding in other aspects. Rapidly reporting bioanalytical results in clinical settings will facilitate medical diagnosis and enable personalized medicine. Our method allows rapid quantification of carboxylic acid metabolites with 5 μL plasma (typically in less than 40 minutes) and requires operation of fewer than 5 minutes per sample, with room for future optimizations. Thanks to its boosted sensitivity, DQmB-HA method simplified the whole workflow and the experimental operation, bypassing vacuum concentration and re-dissolution steps in traditional LC-MS methods for TCA metabolites detection. Therefore, the DQmB protocol is well-matched to automation which promises next-generation high-throughput screening technology based on metabolomics.

## Data availability

The data supporting the findings of this study are available within the paper and its Supplementary Information. Additional information and files are available from the corresponding author upon reasonable request.

## Acknowledgements

This work was supported by Peking-Tsinghua Center for Life Sciences and start-up fund from Peking University (to Z. C.), National Science Foundation of China (82104259 to Q. Z.), and the Special Research Fund for Central Universities, Peking Union Medical College (3332021041 to Q. Z.). We thank Xianxiao Meng and Yanlong Li at the Laboratory Animal Center at Peking University for their technical assistance. We thank Jiaming Mo, Wen Zheng, Yanyun He, and Xiaohong Peng for their technical assistance. All figures in this paper were exported with Adobe Illustrator as vector graphics. Figure 2a, 3a, 4a, 6a, and Supplementary Figure 11 were created with elements from BioRender.

## Author contributions

Z.C. and L.Q. conceived the study and supervised the project. Z.C. and C.L. designed the chemical structure of DQmB. C.L. performed the chemical synthesis. K.C. and C.L. optimized derivatization conditions for DQmB-HA method. Z.C., L.Q. and Q.Z.designed all the experiments involving biomedical samples. Xuelian W. prepared samples containing mouse oocytes and zygotes under the supervision of X.H. Q.Z. conducted isotope labeling experiments with the help from X.H. and K.C., under the supervision of P.L. L.J. performed IR surgeries under the supervision of X.H. Q.Z. and C.L. performed fasting-refeeding experiments with the help from Xiao W., J.L. and D.H. under the supervision of X.C. T.L., X.Z., C.L. and Q.Z. collected blood samples from MELAS induividuals and healthy volunteers under the supervision of Z.W. K.C. and X.L. acquired all the LC-MS/MS data. C.L. and Q.Z. analyzed the LC-MS/MS data under the supervision of L.Q. C.L., L.Q. and Z.C. wrote the manuscript with input from all the authors.

