## Supplementary Information for "Diazo-carboxyl click chemistry enables rapid and sensitive quantification of carboxylic acid metabolites"

### Figures

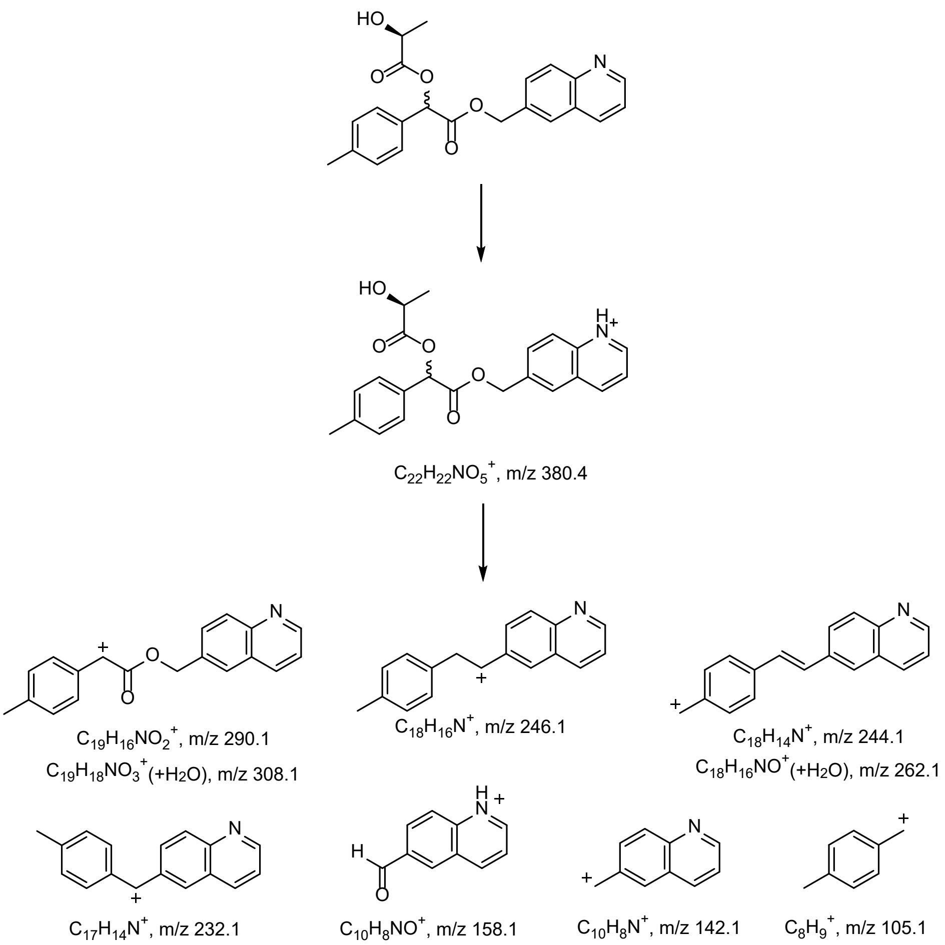

**Supplementary Figure 1**

Typical fragment ions of L-lactate after DQmB-HA derivatization in multiple-reaction-monitoring (MRM) mode.

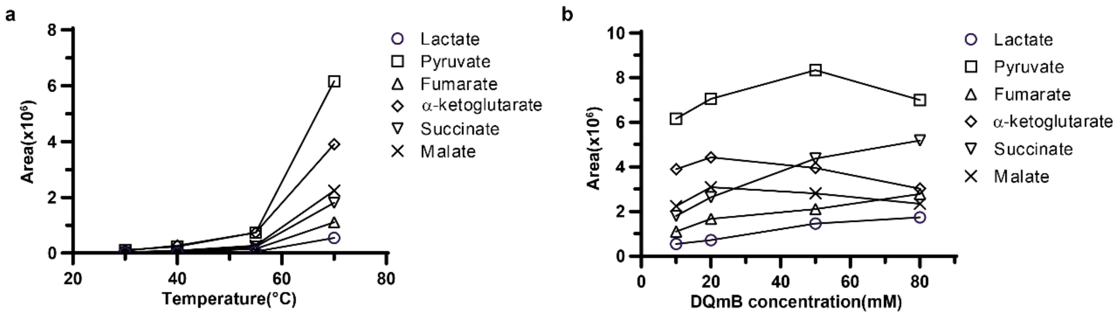

**Supplementary Figure 2**

a,b, Effects of temperature and DQmB concentration on derivatization of carboxylic acids.

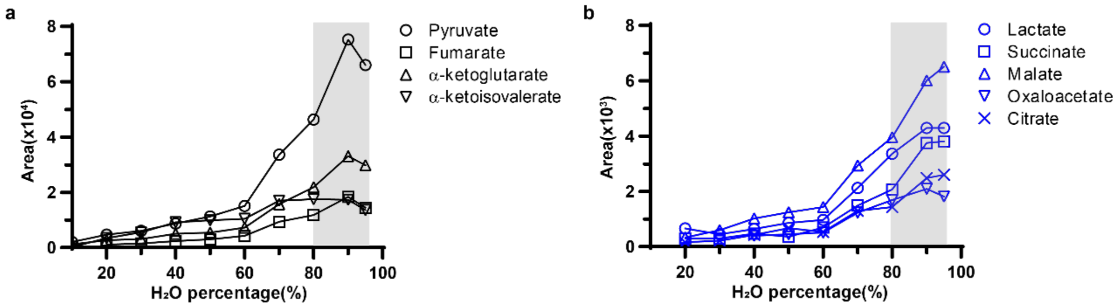

**Supplementary Figure 3**

a,b, Effects of $H_2O$ content(% in volume) on derivatization of carboxylic acid metabolites. Grey shades represent preferred ranges of $H_2O$ content for DQmB-HA derivatization.

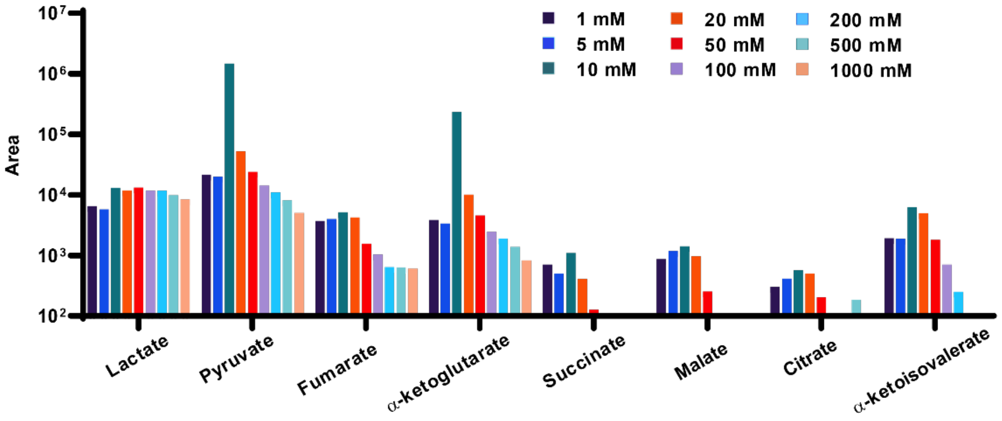

**Supplementary Figure 4**

Effects of hydrochloric acid (HCl) concentration (1-1000 mM) on derivatization of carboxylic acid metabolites.

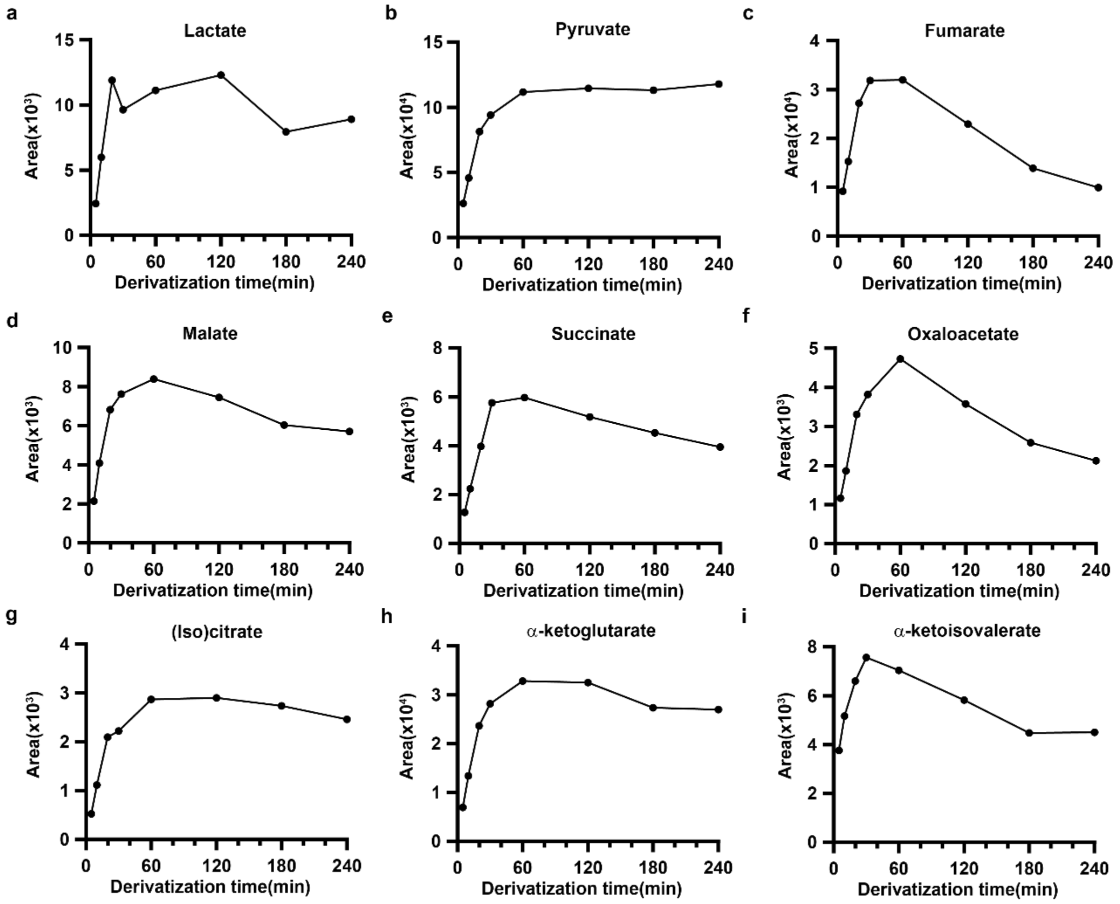

**Supplementary Figure 5**

a-i, DQmB-HA derivatization kinetics of lactate (a), pyruvate (b), fumarate (c), malate (d), succinate (e), oxaloacetate (f), (iso)citrate (g), α-ketoglutarate (h), α-ketoisovalerate (i). Derivatization setup: 70 °C, 10 μL mixed standard solution (1 μM for each carboxylic acid), 40 μL solution A, 50 μL solution B.

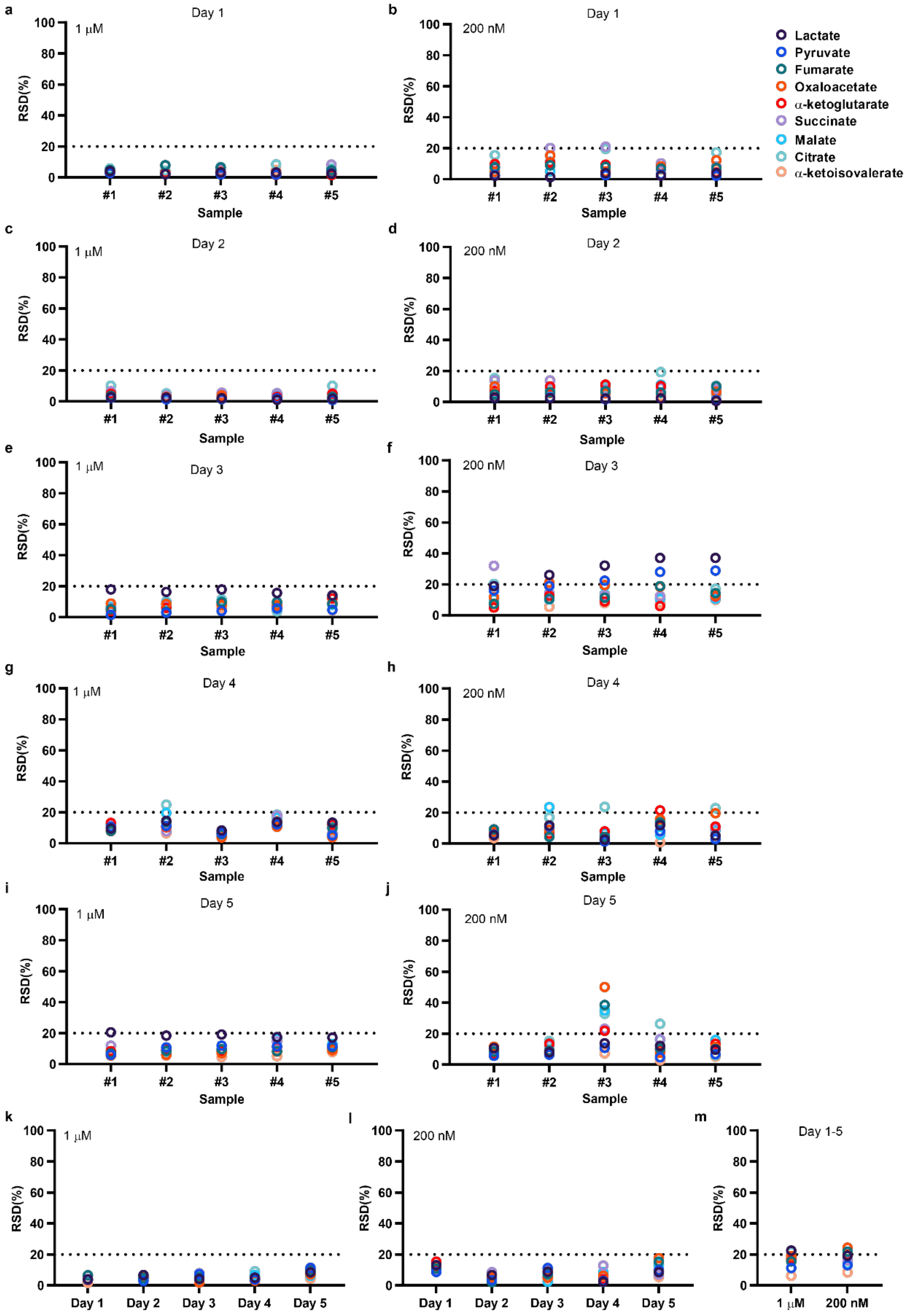

**Supplementary Figure 6**

a-j, Intraday technical repeatability of derivatized carboxylic acids (1 μM and 200 nM) in Day 1(a,b), Day 2(c,d), Day 3(e,f), Day 4(g,h), Day 5(i,j). Relative standard deviations (RSDs) were calculated from five injections for each sample on Day 1, 2, 3, and 5, and from three injections for each sample on Day 4.

k, l, Intraday sample repeatability of carboxylic acids (1 μM and 200 nM) derivatized

by DQmB-HA method. RSDs were calculated from five samples on each day.
m, Interday repeatability of carboxylic acids (1 μM and 200 nM) derivatized by DQmB-HA method. RSDs were calculated from the daily average MS intensities of five days.

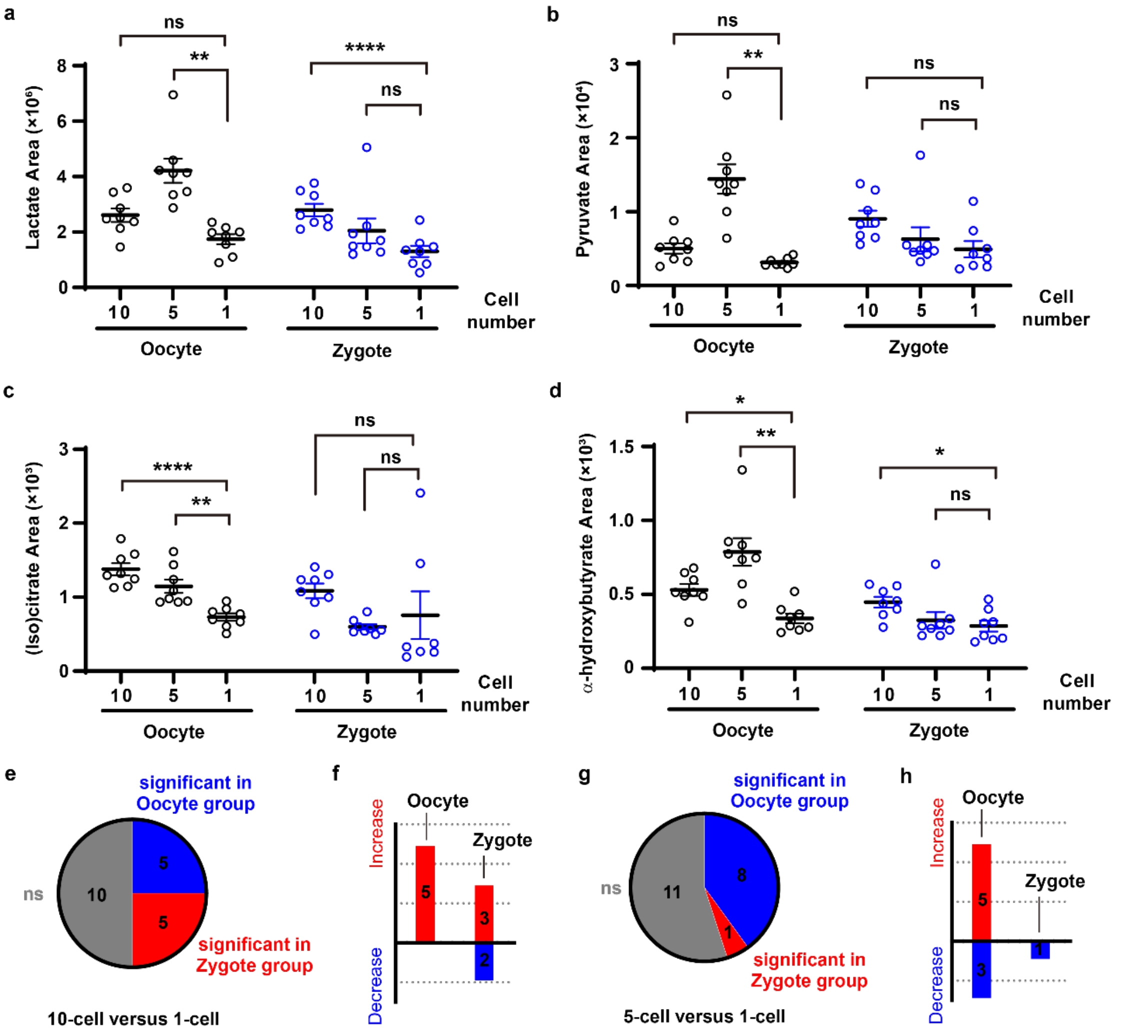

**Supplementary Figure 7**

a-d, Measured levels of lactate (a), pyruvate (b), (iso)citrate (c), and α-hydroxybutyrate (d) in 1-cell, 5-cell, and 10-cell samples separately. Data are mean ± s.e.m.; one-way ANOVA followed by selected multiple comparisons (n=8 for each group, 10-cell versus 1-cell and 5-cell versus 1-cell as selected pairs).

e, g, Pie charts showing significance distribution of comparing 10-cell sample versus 1-cell sample (e), or 5-cell sample versus 1-cell sample (g) in Oocyte group and Zygote group.

f, h, The directions of change in 10-cell samples (f) or 5-cell samples (h) compared to 1-cell samples.

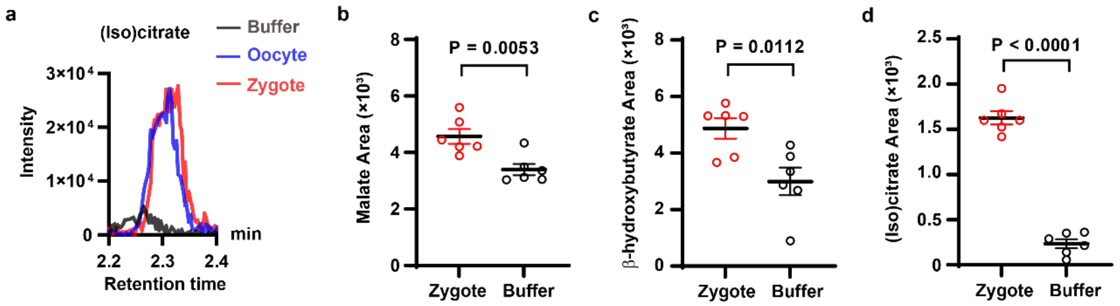

**Supplementary Figure 8**

a, Representative chromatograms of (iso)citrate in buffer, 10 oocytes, or 10 zygotes. b-d, Comparisons of malate (b), β-hydroxybutyrate (c), (iso)citrate (d) in 10 zygotes and buffer. Data are mean ± s.e.m.; unpaired Student's *t* test (n=6 for each group).

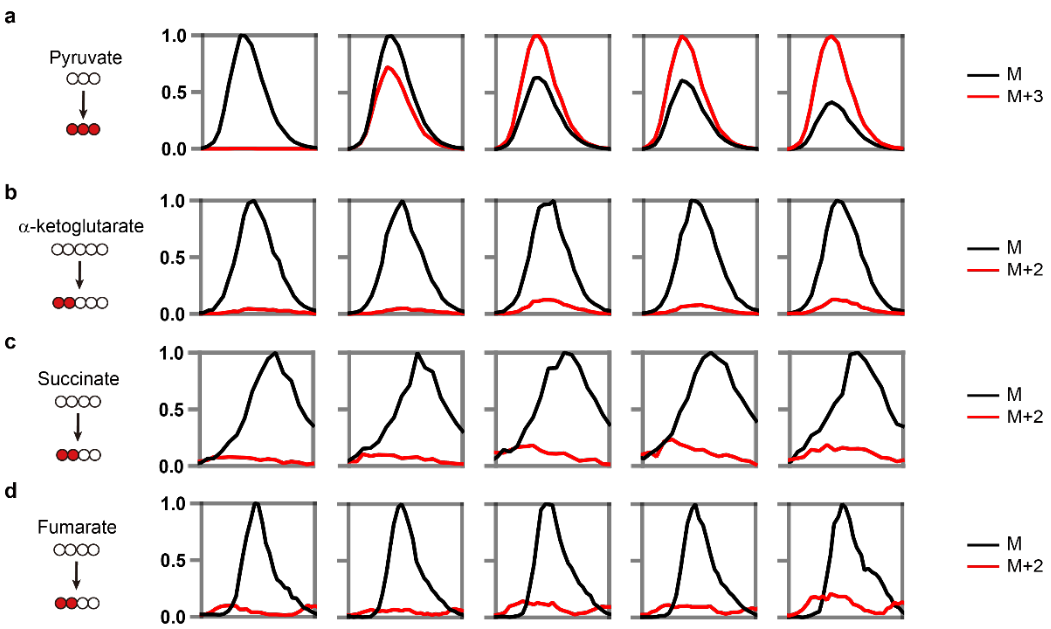

**Supplementary Figure 9**

a-d, Representative merged chromatograms showing both labeled and unlabeled fractions of pyruvate (a), α-ketoglutarate (b), succinate (c), and fumarate (d) at multiple time points in control NRVMs.

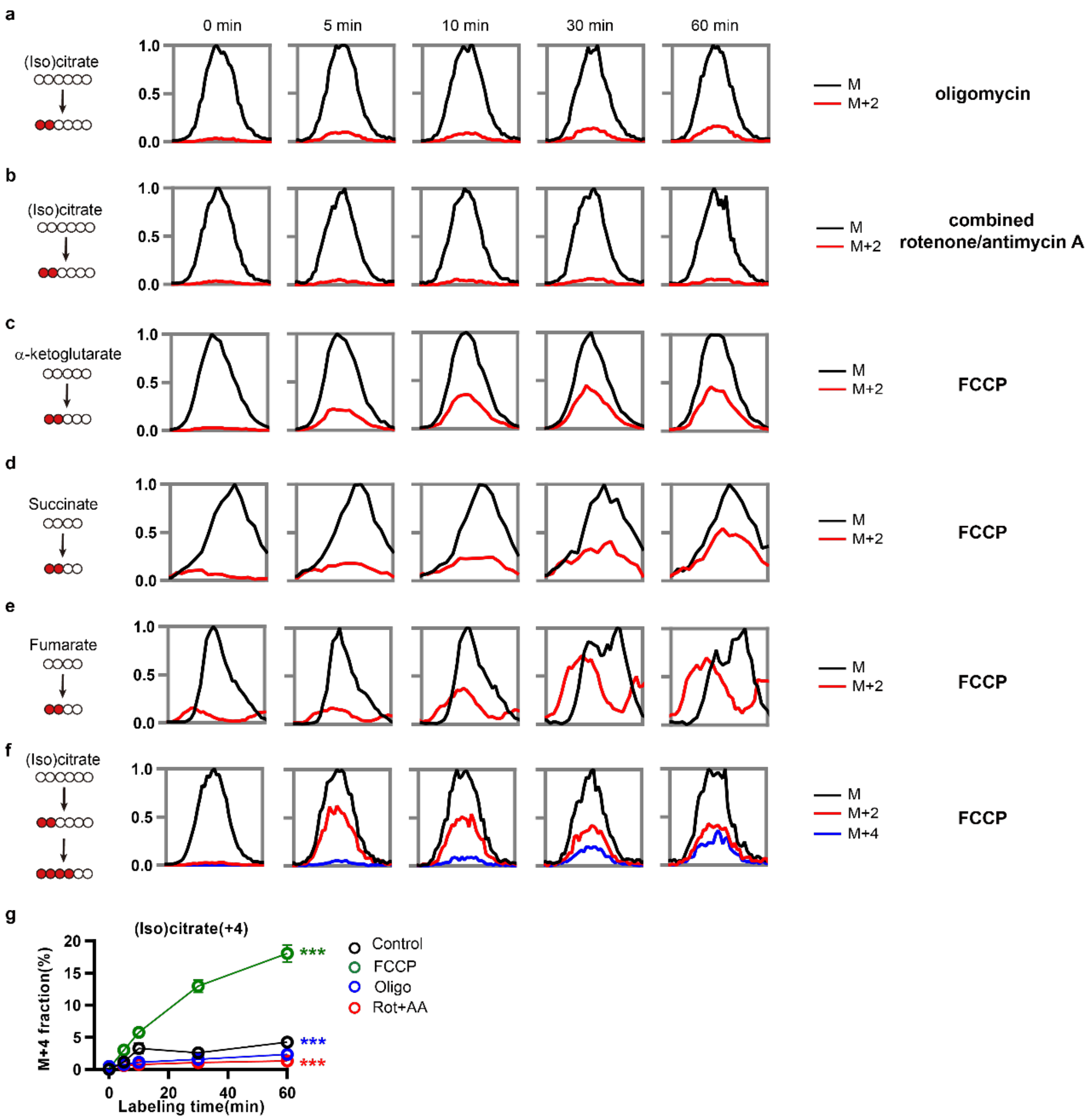

**Supplementary Figure 10**

a, b, Representative merged chromatograms showing both labeled and unlabeled fractions of (iso)citrate at multiple time points in NRVMs treated with oligomycin or combined rotenone/antimycin A.

c-f, Representative merged chromatograms showing both labeled and unlabeled fractions of α-ketoglutarate (c), succinate (d), fumarate (e), and (iso)citrate (f), at multiple time points in NRVMs treated with FCCP.

g, Comparisons of the labeled fractions of (iso)citrate (M+4) at 0, 5, 10, 30, and 60 minutes in NRVMs treated with FCCP, Oligo, or Rot+AA separately, to control. Data are mean ± s.e.m.; one-way ANOVA followed by multiple comparisons (n=8 for each group).

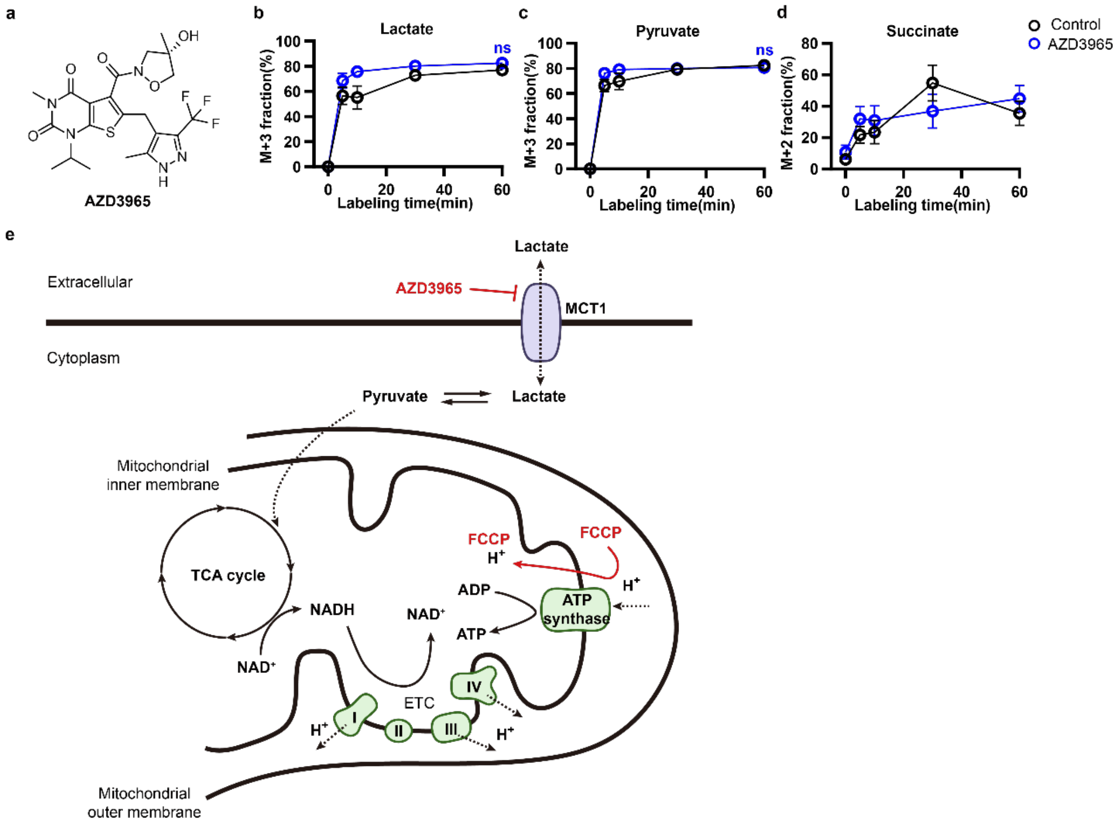

**Supplementary Figure 11**

a, Chemical structure of AZD3965.

b-d, Comparisons of the labeled fractions of lactate (b), pyruvate (c), and succinate (d) at 0, 5, 10, 30, and 60 minutes in control NRVMs and AZD3965-treated NRVMs. Data are mean ± s.e.m.; unpaired Student's *t* test (n=8 for each group).

e, Working mechanisms of AZD3965 and FCCP.

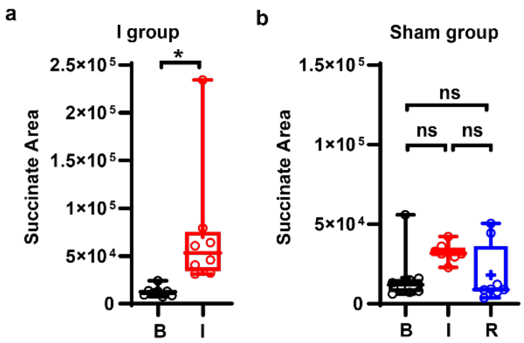

**Supplementary Figure 12**

a, Comparison of the measured succinate levels in mouse serum at Basal (B) and at Ischemia (I) in I group. Data are mean ± s.e.m.; unpaired Student's *t* test (n=8 for each group).

b, Comparisons of the measured succinate levels in mouse serum at Basal (B), Ischemia (I), and Reperfusion (R) in Sham group. Data are mean ± s.e.m.; repeated measures one-way ANOVA followed by multiple comparisons (assume sphericity, n=8 for each group).

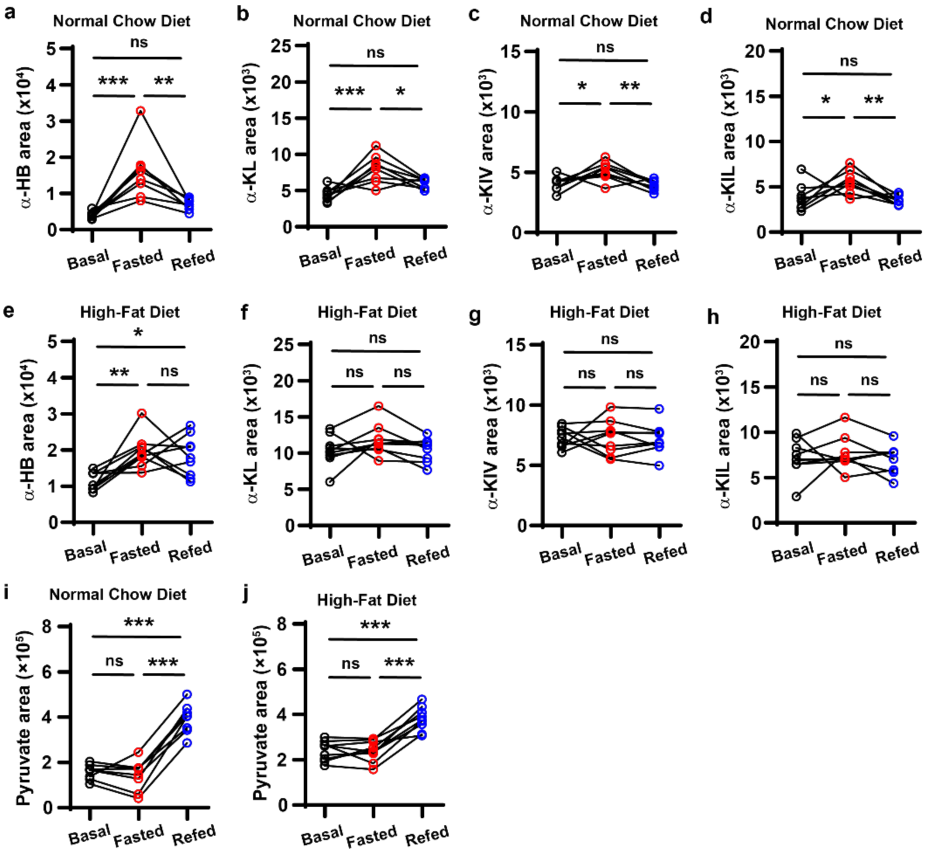

**Supplementary Figure 13**

a-h, Simultaneously monitoring the dynamics of α-hydroxybutyrate, α-ketoleucine, α-ketoisovalerate, and α-ketoleucine before fasting (Basal), after fasting (Fasted), after refeeding (Refed) in serum from NC mice (a-d) and HFD mice (e-h).

i, j, Comparisons of pyruvate before fasting (Basal), after fasting (Fasted), and after refeeding (Refed) in serum from NC mice (i) and HFD mice (j).

Data are mean ± s.e.m.; repeated measures one-way ANOVA followed by multiple comparisons (assume sphericity, n=8 for NC group, n=9 for HFD group).

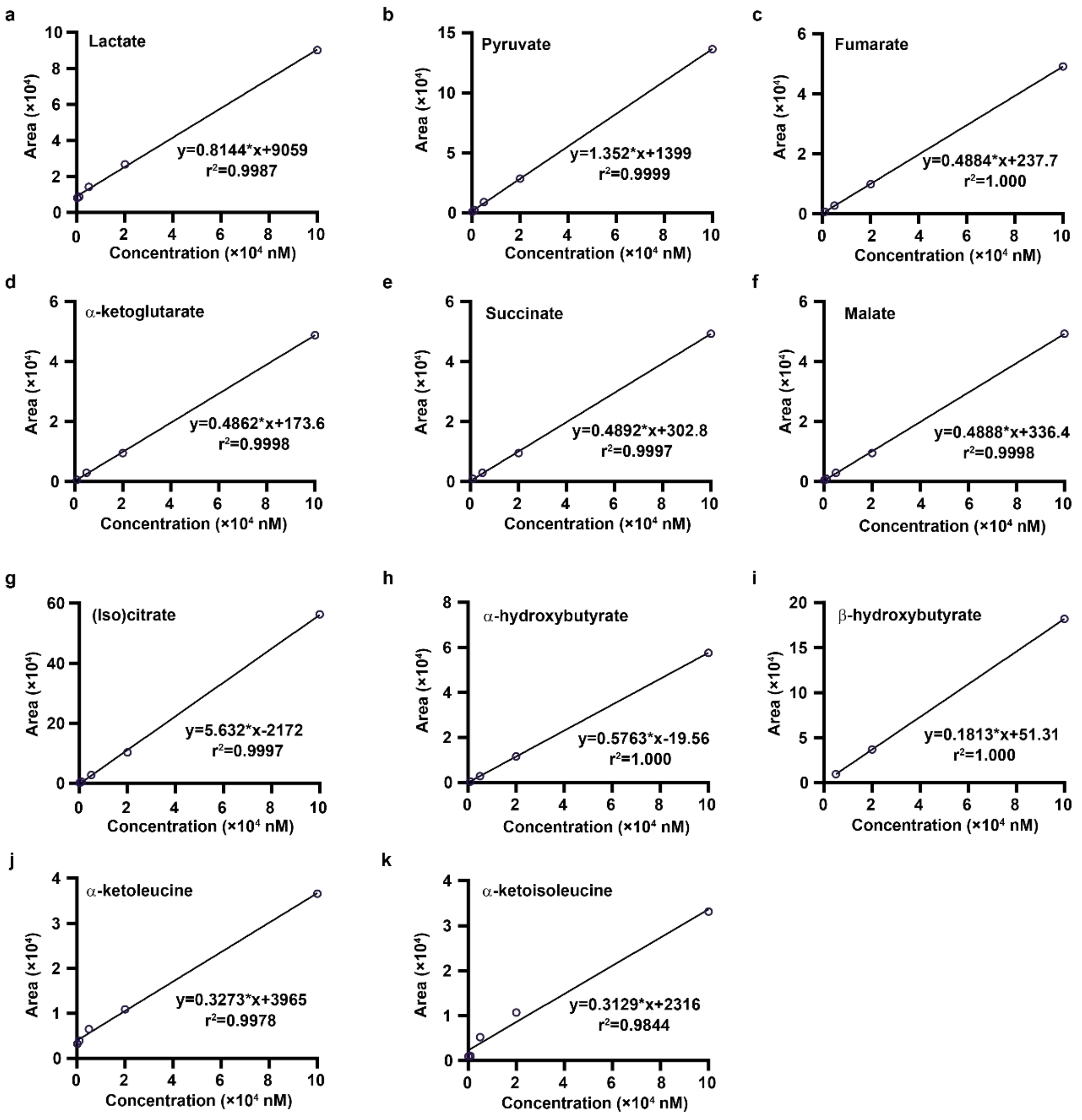

**Supplementary Figure 14**

a-k, external standard curves for the absolute concentration quantification of lactate (a), pyruvate (b), fumarate (c), α-ketoglutarate (d), succinate (e), malate (f), (iso)citrate (g), α-hydroxybutyrate (h), β-hydroxybutyrate (i), α-ketoleucine (j), and α-ketoisoleucine (k).

### Tables

**Table S1 Mass spectrometric parameters for derivatized carboxylic acid metabolites**

| Carboxylic acids | Derivatized by | | Precursor ions(m/z) | Product ions(m/z) | Cone voltage(V) | Collision energy(V) |
|---|---|---|---|---|---|---|
| | DQmB | HA | | | | |
| Lactate | + | - | 380.2 | 142.1 | 28 | 28 |
| Pyruvate | + | + | 393.3 | 142.1 | 22 | 28 |
| (Iso)citrate | + | - | 482.3 | 142.1 | 34 | 34 |
| α-ketoglutarate | + | + | 451.3 | 142.1 | 40 | 40 |
| Succinate | + | - | 408.2 | 142.1 | 28 | 34 |
| Fumarate | + | - | 406.1 | 142.1 | 40 | 28 |
| Malate | + | - | 424.1 | 142.1 | 34 | 34 |
| Oxaloacetate | + | + | 437.1 | 142.1 | 28 | 46 |
| β-hydroxybutyrate | + | - | 394.1 | 141.9 | 40 | 22 |
| α-hydroxybutyrate | + | - | 394.1 | 158.0 | 22 | 40 |
| α-ketoisovalerate | + | + | 421.2 | 232.1 | 40 | 16 |
| α-ketoleucine | + | - | 435.2 | 142.1 | 40 | 40 |
| α-ketoisoleucine | + | - | 435.2 | 142.1 | 40 | 40 |

**Table S2. Signal-to-noise ratios of carboxylic acid metabolites derivatized by DQmB-HA method and oBHA method[1]**

| Metabolites[a] | DQmB-HA method[b] | | oBHA method[c] | | No derivatization[d] | |
|---|---|---|---|---|---|---|
| | Concentration[e] | S/N[f] | Concentration[e] | S/N[f] | Concentration[e] | S/N[f] |
| Lactate | 5 nM | 16.8 | 5 nM | 4.6 | 10 μM | 4.9 |
| Pyruvate | 1 nM | 17.0 | 5 nM | 9.0 | 5 μM | 4.0 |
| Citrate | 10 nM | 1.1 | 5 nM | 2.8 | 0.5 μM | 26.4 |
| α-ketoglutarate | 1 nM | 43.5 | 5 nM | 4.3 | 0.5 μM | 10.4 |
| Succinate | 5 nM | 8.3 | 5 nM | 4.4 | 0.5 μM | 4.6 |
| Fumarate | 0.5 nM | 3.8 | 5 nM | ~134 | 1 μM | 4.0 |
| Malate | 2 nM | 4.4 | 10 nM | 2.5 | 0.5 μM | 26.7 |
| Oxaloacetate | 2 nM | 2.6 | 5 nM | 3.9 | 2 μM | 5.7 |

[a]: Metabolites in acid form were separately dissolved in $H_2O$ as stock solutions of the same concentration. Then, the same volumes from each stock solution were mixed and diluted with $H_2O$ to prepare a series of standard solutions (STD).

[b]: Derivatization: 5 μL STD + 20 μL solution A + 50 μL solution B, 70 °C, 20 min. Injection volume: 1 μL. MS mode: positive.

[c]: Derivatization: 100 μL STD + 50 μL *o*-benzylhydroxylamine (oBHA) in pyridium buffer + 50 μL EDC in pyridium buffer, room temperature, 60 min. Exraction: 300 μL ethyl acetate ×2, dried at 40 °C. Reconstitution: 1 mL 50% methanol/water. Injection volume: 2 μL. MS mode: positive.

[d]: 50 μL STD. Injection volume: 2 μL. MS mode: negative.

[e]: The concentration of each metabolite in STD.

[f]: Signal-to-noise ratios (S/N) were calculated using MassLynx 4.1 software.

**Table S3 P-values[a] of different metabolites in three independent experiments comparing 10 zygotes and 10 oocytes**

| Metabolites | p-value | | |
|---|---|---|---|
| | Independent experiment #1[b] | Independent experiment #2[c] | Independent experiment #3[d] |
| Lactate | 0.3523 | **0.0004** | 0.5955 |
| Pyruvate | 0.2434 | **<0.0001** | **0.0069** |
| Fumarate | 0.2097 | **0.0002** | **<0.0001** |
| Malate | **0.0464** | **0.0011** | **<0.0001** |
| Succinate | 0.2794 | 0.6367 | 0.7917 |
| (Iso)citrate | 0.2370 | 0.1113 | **0.0409** |
| α-ketoleucine | 0.1280 | 0.3266 | 0.5685 |
| α-ketoisoleucine | **0.0056** | 0.5695 | **<0.0001** |
| α-hydroxybutyrate | 0.2007 | **<0.0001** | 0.1466 |
| β-hydroxybutyrate | - | **0.0136** | **0.0012** |

[a]: Unpaired Student's *t* test, two tailed.
[b]: n=3 for Oocyte group, n=3 for Zygote group.
[c]: n=8 for Oocyte group, n=6 for Zygote group.
[d]: n=8 for Oocyte group, n=8 for Zygote group.

#### Table S4 Information of MELAS individuals

| Case number | | 3841 | 3822 | 3859 | 3853 | 3815 | 3942 | 4477 | 4500 | 5353 |
|---|---|---|---|---|---|---|---|---|---|---|
| Age(years) | | 26 | 25 | 29 | 18 | 34 | 42 | 19 | 20 | 36 |
| Gender | | F | M | M | M | M | F | M | F | F |
| Disease duration(years) | | 6 | 1 | 1 | 3 | 13 | 12 | 2 | 6 | 1 |
| Genotype | | m. 3243A>G | m. 3243A>G | m. 3243A>G | m. 3243A>G | m. 3243A>G | m. 13513G>A | m. 3243A>G | m. 3243A>G | m. 3243A>G |
| Clinical symptoms | Abnormal behavior | - | - | - | - | - | - | + | - | - |
| | Aphasia | - | - | - | - | - | - | - | - | - |
| | Cognitive impairment | + | - | - | - | + | - | - | - | - |
| | Cortical blindness | - | + | - | - | - | - | + | - | + |
| | Exercise intolerance | + | - | - | - | + | - | - | + | - |
| | Headache | + | + | - | - | - | + | + | - | + |
| | Hemiplegia | - | - | - | - | - | - | - | - | - |
| | Hyperthermia | - | + | - | - | - | - | - | - | - |
| | Seizures | - | + | + | + | + | + | - | + | + |
| | Visual abnormality | + | - | - | - | + | - | + | - | - |
| | Vomiting | + | - | - | - | - | - | - | + | + |

**Table S5 Information of healthy volunteers**

| ID | 01 | 02 | 03 | 04 | 05 | 06 | 07 | 08 | 09 | 11 | 13 |
|---|---|---|---|---|---|---|---|---|---|---|---|
| Age(years) | 27 | 27 | 22 | 24 | 22 | 25 | 25 | 24 | 23 | 23 | 37 |
| Gender | M | M | M | M | M | F | F | F | F | M | F |

**Table S6. Parameters for principle component analysis (PCA) of healthy volunteers and MELAS individuals**

| Metabolites | Loadings | |
|---|---|---|
| | PC1 | PC2 |
| Fumarate | 0.3972 | 0.2086 |
| Malate | 0.3914 | 0.2644 |
| Succinate | 0.3783 | 0.2140 |
| Lactate | 0.3744 | 0.2910 |
| α-hydroxybutyrate | 0.3347 | -0.1520 |
| Pyruvate | 0.2804 | 0.0541 |
| β-hydroxybutyrate | 0.2521 | -0.2891 |
| α-ketoisoleucine | 0.2492 | -0.5449 |
| α-ketoleucine | 0.2345 | -0.5772 |
| α-ketoglutarate | 0.1747 | -0.0245 |
| (Iso)citrate | -0.0623 | -0.1258 |

### Experimental Procedures

#### DQmB-HA derivatization

Preparations of solution A1, A2, A and B. Solution A1 contains 20 mM DQmB in acetonitrile. Solution A2 contains 20 mM HA(HCl salt) in $H_2O$. Solution A is prepared from solution A1 and solution A2 (v/v=20:1), containing ~20 mM DQmB and ~1 mM HA. Solution B contains 10 mM hydrochloric acid in $H_2O$.

For carboxylic acid standards, 5 μL standard solution was added with solution A (20 μL) and solution B (50 μL) and vortexed for 30 seconds. Then the mixture was incubated at 70 °C for 20 min, and centrifuged at 14,000 rpm (Eppendorf 5424R), 4 °C for 10 min. The supernatant solution was used for LC-MS/MS analysis.

For oocytes and zygotes, solution A (5 μL) and solution B (10 μL) were immediately added into PCR tubes sequentially. The mixture was heated at 70 °C for 20 min and centrifuged at 14,000 rpm (Eppendorf 5424R), 4 °C for 10 min. 5 μL supernatant solution was used for LC-MS/MS analysis.

For isotope-labeled NRVMs, solution A (25 μL) was added to each well after rinsing and the whole 96-well plate was sonicated for 30 seconds in a water bath. Then solution B (50 μL) was added and the mixture in each well was transferred to 200 μL Eppendorf tubes and heated at 70 °C for 20 min. The derivatized samples were centrifuged at 14,000 rpm (Eppendorf 5424R), 4 °C for 10 min, and the supernatant solution was used for LC-MS/MS analysis.

For mouse plasma and human plasma, 5 μL plasma was added with solution A (20 μL) and solution B (50 μL) and vortexed for 30 seconds. Then the mixture was incubated at 70 °C for 20 min, and centrifuged at 14,000 rpm (Eppendorf 5424R), 4 °C for 10 min. The supernatant solution was used for LC-MS/MS analysis.

**LC-MS/MS**

Liquid chromatography-tandem mass spectrometry (LC-MS/MS) was performed on a UPLC-MS/MS system (I-Class TQ-S micro, Waters, Milford, MA). Chromatographic separation was performed on a Cortecs HSS T3 column (1.6 µm, 2.1×100 mm, Waters, Milford, MA). Gradient elution was applied at a flow rate of 0.4 mL/min and the column thermostat was set at 40 °C. Mobile phases A and B were water and acetonitrile supplemented with 0.1% formic acid (v/v), respectively. Initial elution started with 25% B, and at 0.5 min a gradient (curve 11) was started from 25% to 75% B in 4.5 min. In the following 0.1 min, the percentage of mobile phase B was increased to 100% and maintained for 2 min for column wash. The elute composition returned to the initial conditions in 0.1 min, and the column was equilibrated at the initial conditions for 1 min before the next injection, leading to a total run time of 8 min. The auto-sampler temperature was set at 4 °C. Unless mentioned otherwise, the injection volume was 2 µL. The same chromatographic conditions were used throughout this work.

For mass spectrometry, electrospray ionization (ESI) source in positive mode or negative mode was used. The source temperature was set at 150 °C. The solvent was deprived at a 1000 L/hr high-purity nitrogen flow at 400 °C. The ionization was conducted at a 2 kV capillary voltage. The sampling cone was equipped with 30 kV and a 20 L/hr high-purity nitrogen flow. Multi-reaction monitoring (MRM) mode was used to reduce the matrix effect, of which the parameters were listed in Table S1. All the data acquisition and processing were performed by MassLynx 4.1 software.

**Animals**

Animals were maintained in the Laboratory Animal Center at Peking University, Beijing, China. This facility is accredited by AAALAC. All procedures were performed following protocols approved by the IACUC of Peking University, and conformed to the Guide for the Care and Use of Laboratory Animals (NIH publication No. 86-23, revised 2011).

**Collection of mouse oocytes and zygotes**

Wild-type C57BL/6J female mice (4-6 weeks) were super-ovulated by intraperitoneal injection of PMSG (10 IU/each) and hCG (10 IU/each) with a 48-h interval. Then, the super-ovulated female mice were randomly divided into two groups (Unmated Group and Mated Group). For Mated Group, female mice were immediately mated one-to-one with wild-type C57BL/6J male mice.

All female mice were euthanized with $CO_2$ at 18 h post hCG injection. Zygotes or oocytes were dissected out of the ampulla in the oviduct. The embryo cumulus complexes were treated with 300 μg/ml of hyaluronidase for less than 1 min to disperse the cumulus cells, washed in modified KSOM medium without pyruvate/glucose, and transferred to M2 medium (Millipore, MR-015-D). Then, 1, 5, or 10 zygotes or oocytes were picked with a microcapillary under a dissection microscope, washed in PBS, and transferred into 200 μL polymerase chain reaction (PCR) tubes (Axygen, PCR-0208-C) for DQmB-HA derivatization.

**Isotope labeling of cells in 96-well plates**

The isolation of neonatal rat ventricular myocytes (NRVMs) was performed as described previously[2–4]. Briefly, hearts from 1-day-old Sprague-Dawley rats were removed, and the ventricles were separated from attached vessels and atria and then trisected. NRVMs were prepared from digestion of these tissue fragments.

Isolated NRVMs were plated at a density of $\sim 1\times10^5$ cells/well in 96-well plates and cultured in Dulbecco's modified Eagle's medium (DMEM, Invitrogen) supplemented with 10% fetal bovine serum (FBS, Gibco) in the presence of 0.1 mM 5-bromo-2-deoxyuridine (Sigma). About 12 h after initial plating, medium in each well was replaced by glucose-free DMEM for 4 h. Then, medium was again replaced with 50 μL glucose-free DMEM containing DMSO (Control group) or oligomycin (1 μM) or FCCP (5 μM) or combined antimycin A/rotenone (0.5 μM/0.5 μM). After 5-minute incubation, 50 μL DMEM containing DMSO (Control group) or oligomycin (1 μM) or FCCP (5 μM) or combined rotenone/antimycin A (0.5 μM/0.5 μM) supplemented with 40 mM U-$^{13}C_6$-glucose (Cambridge Isotope Laboratories, CLM-1396-PK) were added into each well of corresponding groups and isotope labeling was started. At each time point from 0 min to 60 min, isotope labeling was quenched by the removal of DMEM supplemented with U-$^{13}C_6$-glucose. Then, cells were rinsed with phosphate buffer saline (PBS) to remove residual medium for further DQmB-HA derivatization.

#### Myocardial ischemia-reperfusion (IR) mouse model

Myocardial IR surgeries were performed as previously described[5,6].

Wild-type C57/BL6J male mice (7-10 weeks) were first randomly divided into different groups. For IR group, animals were anesthetized with avertin (2.5%, 0.01 mL/g, i.p.) and ventilated via a tracheostomy on a Minivent respirator. At this time point (Basal), 50 μL blood was collected and stored in 1.5 mL Eppendorf tubes on ice. A midline sternotomy was performed, and a reversible coronary artery snare occluder was placed around the left anterior descending coronary artery. Animals were challenged with cardiac ischemia by tightening the snare occluder for 30 min. At this time point, 50 μL blood was collected and stored in 2 mL Eppendorf tubes on ice. Then reperfusion was initiated by loosening the snare occluder and mice were allowed to recover for 5 min or 24 h. At the end of 5-min or 24-h reperfusion, 50 μL blood was collected and stored in 2 mL Eppendorf tubes on ice. All blood samples were collected from mice tail tips by heparin capillary, and centrifuged at 4 °C for 10 min at 3000 rpm to obtain serum for further DQmB-HA derivatization.

For Ischemic group, the same surgery for IR group was performed except that reperfusion was not performed and 50 μL blood was not collected at the end of 24-h reperfusion. For Sham group, the same surgery for IR group was performed except that the snare occluder was not tightened.

#### Fasting-refeeding mouse model

For high-fat-diet (HFD) group, 2-month-old male mice were fed with a rodent diet with 60% kilocalories from fat (Research Diets, D12492) for 12 weeks. For normal-chow-diet (NC) group, 2-month-old male mice were fed with normal chow diet for 12 weeks.

Immediately before fasting, 50 μL blood was collected. Acute fasting was initiated at 8 p.m. on Day 1. After 12-h fasting, 50 μL blood was collected at 8 a.m. on Day 2. Mice were again fed with high-fat diets or normal chow diets, respectively. After 6-h refeeding, 50 μL blood was collected at 2 p.m. on Day 2. All blood samples were collected from mice tail tips by heparin capillary, and centrifuged at 4 °C for 10 min at 3000 rpm to obtain plasma for further DQmB-HA derivatization.

#### Collection of human blood samples

This study was approved by the Local Ethics Committee of Peking University First Hospital. All participants' consent was obtained according to the Declaration of Helsinki for use of their remaining samples for research after the diagnosis.

Intravenous blood of MELAS individuals and healthy volunteers were collected and centrifuged at 2000 g, 4 °C for 10 min. Plasma or serum were immediately collected from the supernatants and stored at -80 °C until DQmB-HA derivatization.

#### Statistics

Statistical analysis was performed using GraphPad Prism (Graphpad Software) unless otherwise noted. Sample sizes were determined based on previous experience.

No statistical methods were used to predetermine sample size. Data distribution was assumed to be normal, but this was not formally tested. ns: not significant ($p > 0.05$), *: $p < 0.05$, **: $p < 0.01$, ***: $p < 0.001$, ****: $p < 0.0001$. Z scores were calculated by subtracting the population mean from an individual raw score and then dividing the diffenrence by the population standard deviation. PCA was performed using MetaboAnalyst 5.0[7].

**Organic synthesis and characterization**

General methods

Unless otherwise noted, reagents and solvents were obtained from Energy Chemical, J&K Scientific, and Innochem and used without further purification.

Reactions were monitored by thin layer chromatography (TLC) on precoated TLC plates (silica gel 60 $GF_{254}$, 0.2 mm thickness). TLC chromatograms were visualized using UV light (254 nm) or developed with $KMnO_4$ stain. Column chromatography was performed on manually packed silica gel (200-300 mesh, Qingdao Haiyang Chemicals).

Nuclear magnetic resonance (NMR) spectra were recorded on a Bruker 400 ($^{1}$H 400 MHz, $^{13}$C 101 MHz) Fourier Transform (FT) NMR spectrometers. $^{1}$H and $^{13}$C chemical shifts (δ) were referenced to TMS or residual solvent peaks. Data for $^{1}$H NMR spectra are reported as follows: chemical shift (δ ppm), multiplicity (s = singlet, d = doublet, t = triplet, q = quartet, dd = doublet of doublets, dt = triplet of doublets, m = multiplet, br = broad), coupling constant (Hz), integration. Data for $^{13}$C NMR are reported by chemical shift (δ ppm).

Chemical synthesis of DQmB

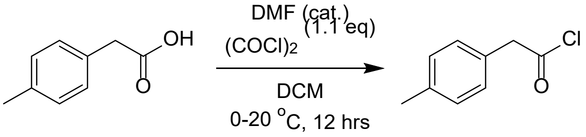

1 2

To a solution of compound **1** (10 g, 66.6 mmol, 1 eq) in anhydrous DCM (60 mL) at 0-5 °C was added oxalyl chloride (9.30 g, 73.3 mmol, 6.41 mL, 1.1 eq) and DMF (48.7 mg, 666 μmol, 51.2 μL, 0.01 eq) at 0 °C. The reaction mixture was stirred at 20 °C for 12 hrs and concentrated under vacuum to yield compound **2** (12 g, crude) as a yellow oil which was used directly in the next step without further purification.

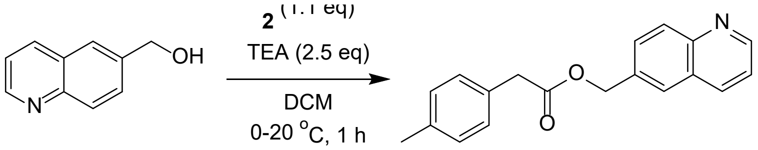

3 4

To a solution of compound **3** (6 g, 37.7 mmol, 1 eq) and TEA (13.1 mL, 94.2 mmol, 2.5 eq) in DCM (36 mL) was added compound **2** (6.99 g, 41.5 mmol, 1.1 eq) at 0 °C. The reaction was stirred at 20 °C for 15 hrs and reaction crude was poured into water (30 mL). The aqueous phase was extracted with DCM twice and the combined organic phase was dried with anhydrous $Na_2SO_4$, filtered, and concentrated in vacuum. The residue was purified by column chromatography on silica gel to yield compound **4** (6.8 g, 23.3 mmol, 62% yield) as a yellow solid. $^{1}$H NMR (400 MHz, $CDCl_3$): δ 8.91 (dd, *J* = 4.2, 1.5 Hz, 1H), 8.11 - 8.06 (m, 2H), 7.69 (s, 1H), 7.63 (dd, *J* = 8.7, 2.0 Hz, 1H), 7.39 (dd, *J* = 8.3, 4.2 Hz, 1H), 7.17 (dd, *J* = 24.0, 8.0 Hz, 4H), 5.30 (s, 2H), 3.67 (s, 2H), 2.34 (s, 3H). $^{13}$C NMR (101 MHz, $CDCl_3$): δ 171.6, 150.8, 148.0, 136.9, 136.1, 134.3, 130.8, 129.9, 129.4, 129.2, 129.2, 128.0, 126.8, 121.5, 66.1, 41.0, 21.1. MS (ESI) calcd for $C_{19}H_{18}NO_2$ $[M+H]^+$ 292.1, found 292.1.

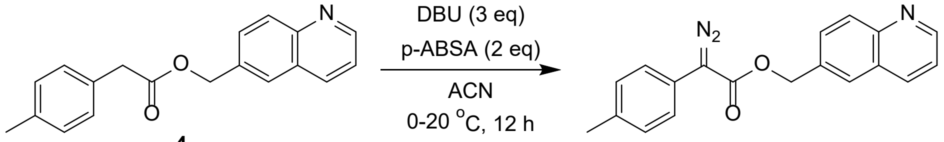

**4** **DQmB**

To a solution of compound **4** (6.8 g, 23.3 mmol, 1 eq) in ACN (45 mL) was added p-ABSA (9.21 g, 46.7 mmol, 2 eq) and DBU (10.7 g, 70.0 mmol, 10.6 mL, 3 eq) at 0 °C. The reaction was stirred at 20 °C for 12 hrs and concentrated under reduced pressure to give a residue. The residue was purified by column chromatography on silica gel to yield **DQmB** (5.8 g, 17.3 mmol, 74% yield) as an orange solid. $^{1}$H NMR (400 MHz, $CDCl_3$): δ 8.93 (dd, *J* = 4.2, 1.7 Hz, 1H), 8.16 (dd, *J* = 8.4, 1.0 Hz, 1H), 8.13 (d, *J* = 8.7 Hz, 1H), 7.83 (d, *J* = 1.1 Hz, 1H), 7.74 (dd, *J* = 8.7, 1.9 Hz, 1H), 7.42 (dd, *J* = 8.3, 4.2 Hz, 1H), 7.37 (d, *J* = 8.3 Hz, 2H), 7.20 (d, *J* = 8.1 Hz, 2H), 5.48 (s, 2H), 2.34 (s, 3H). $^{13}$C NMR (101 MHz, $CDCl_3$): δ 165.2, 150.8, 148.1, 136.1, 135.9, 134.3, 130.1, 129.7, 129.3, 128.0, 127.1, 124.2, 121.9, 121.5, 66.1, 21.0. MS (ESI) calcd for $C_{19}H_{16}N_3O_2$ $[M+H]^+$ 318.1, found 318.1, 290.1 ($-N_2$).

LC-MS chromatograms

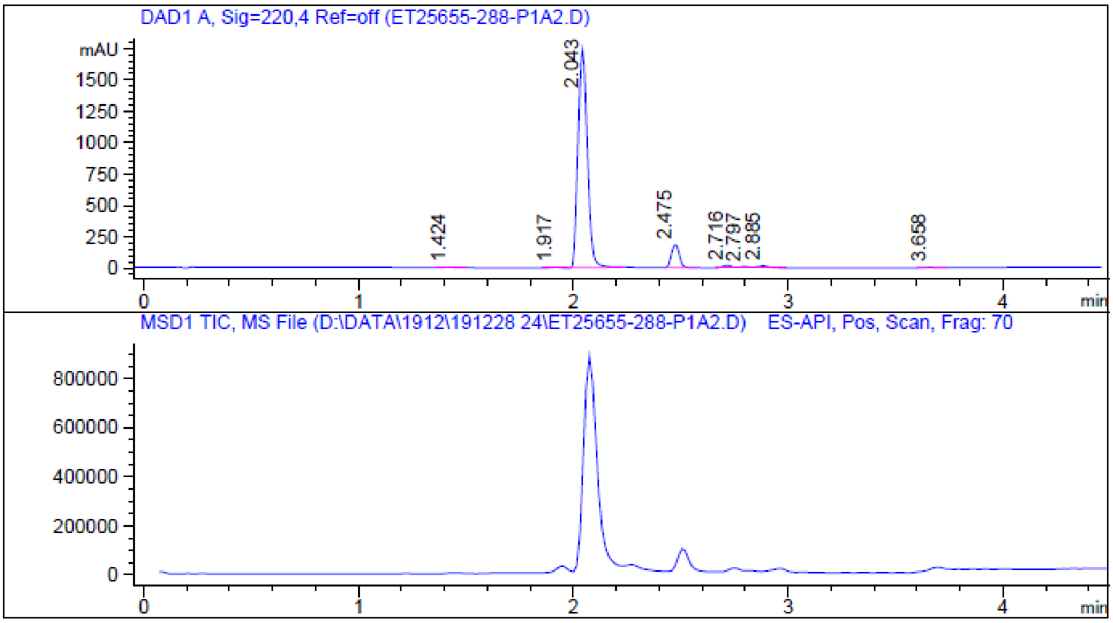

LC-MS chromatograms of **Compound 4**. X-axis: retention time. Y-axis: 220 nm (top), total ion current (TIC, bottom).

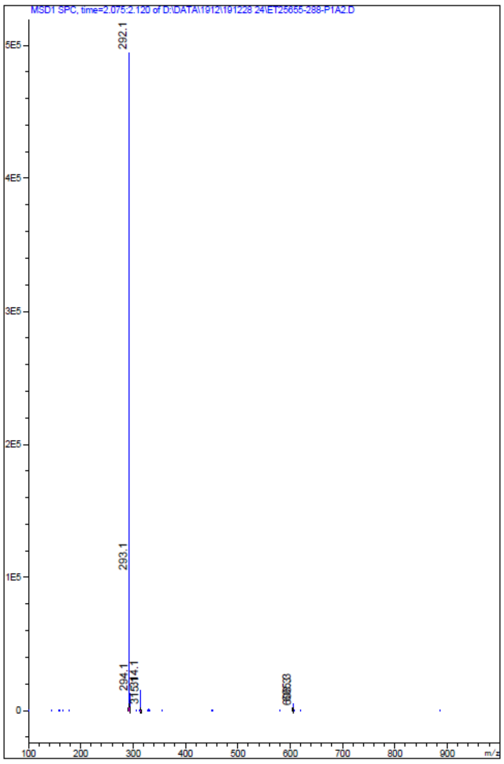

Mass spectrum of the peak eluted at 2.043 min.

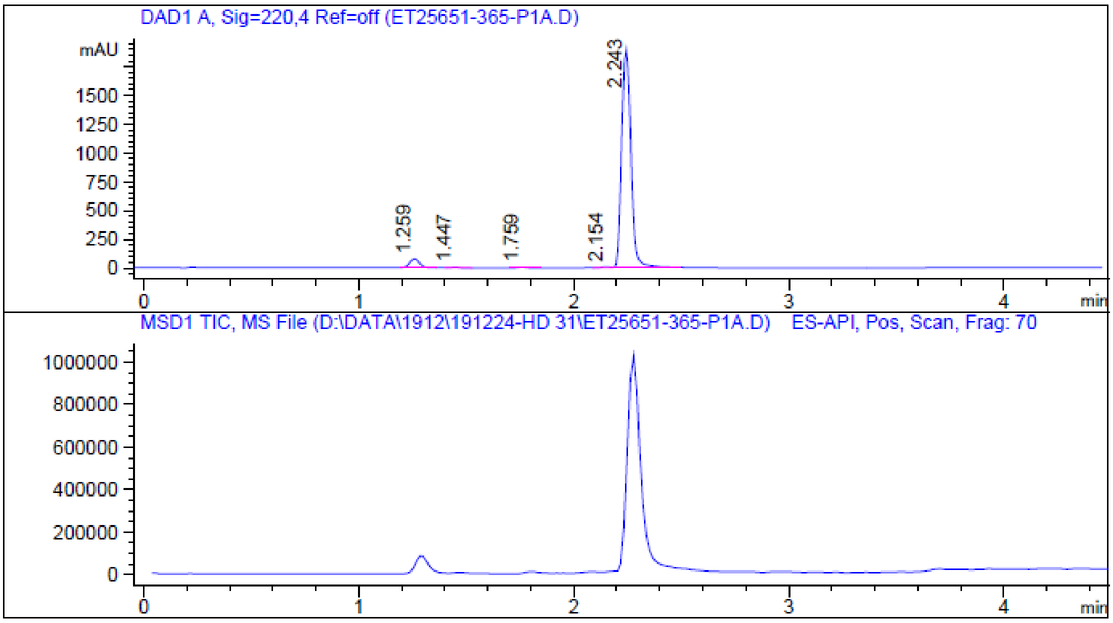

LC-MS chromatograms of **DQmB**. X-axis: retention time. Y-axis: 220 nm (top), total ion current (TIC, bottom).

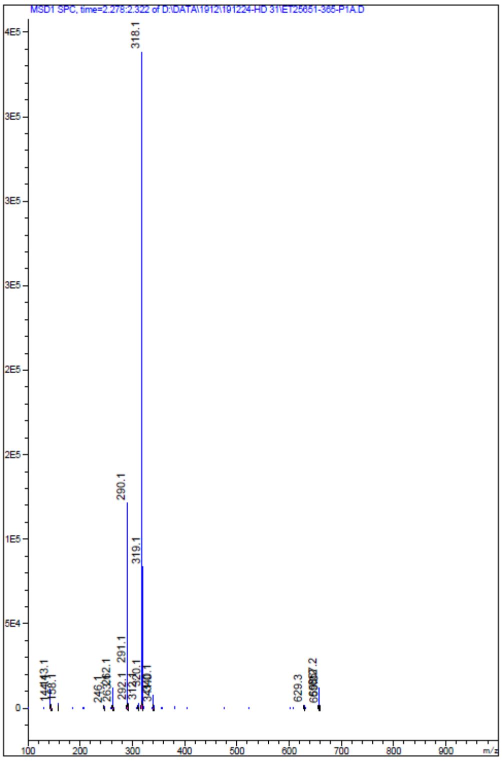

Mass spectrum of the peak eluted at 2.243 min.

NMR spectra

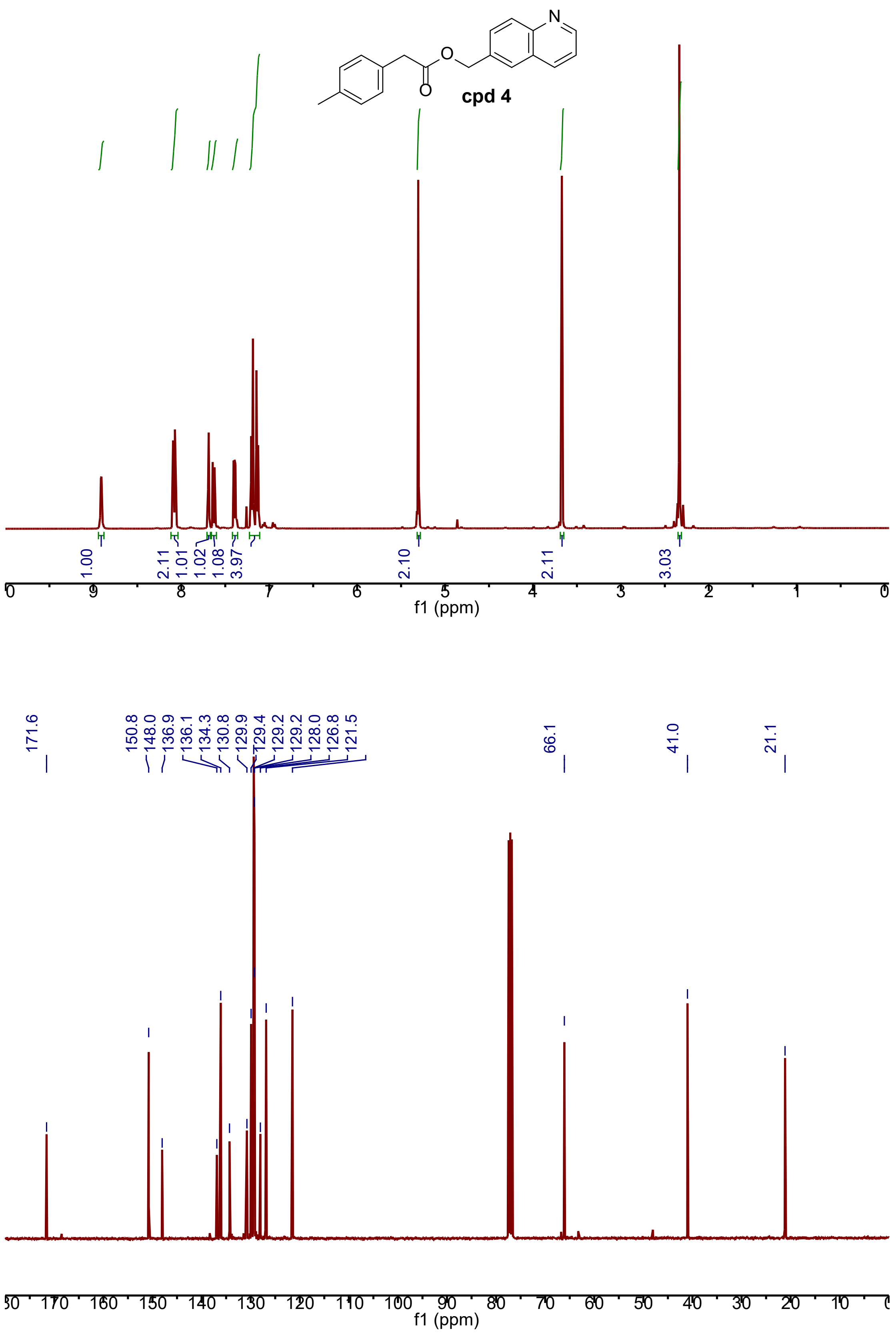

[1]H NMR spectrum(top) and [13]C NMR spectrum(bottom) of **Compound 4**.

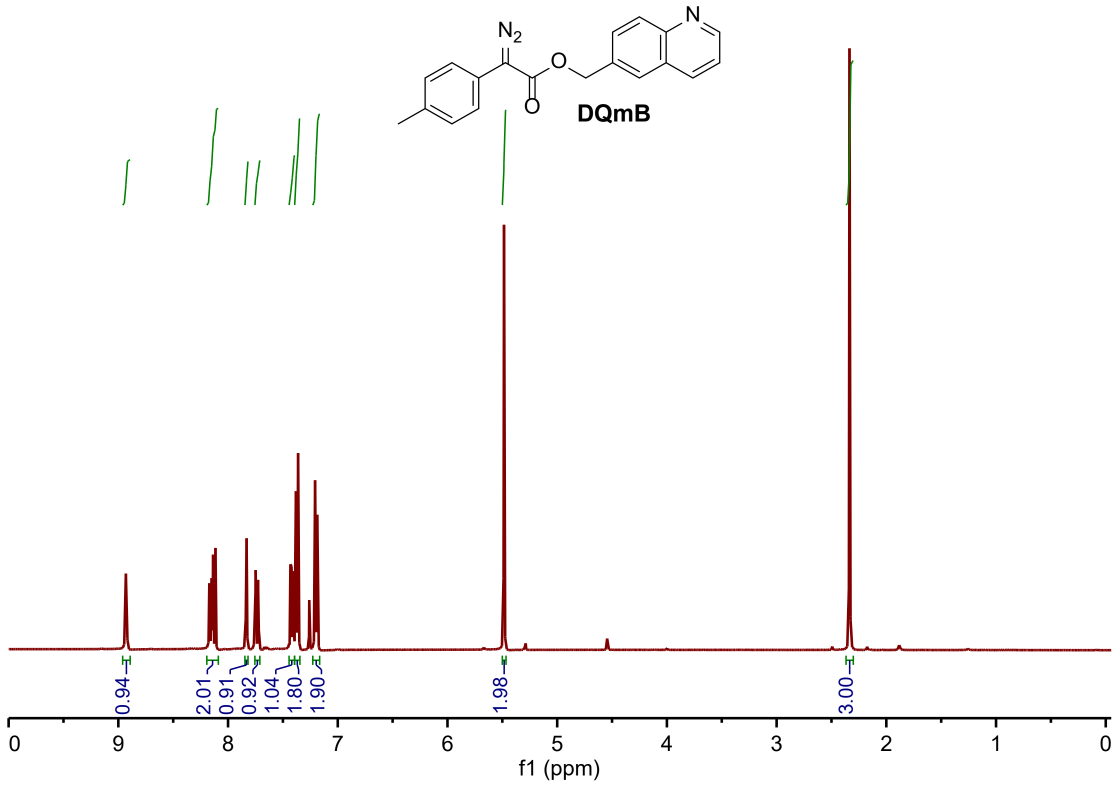

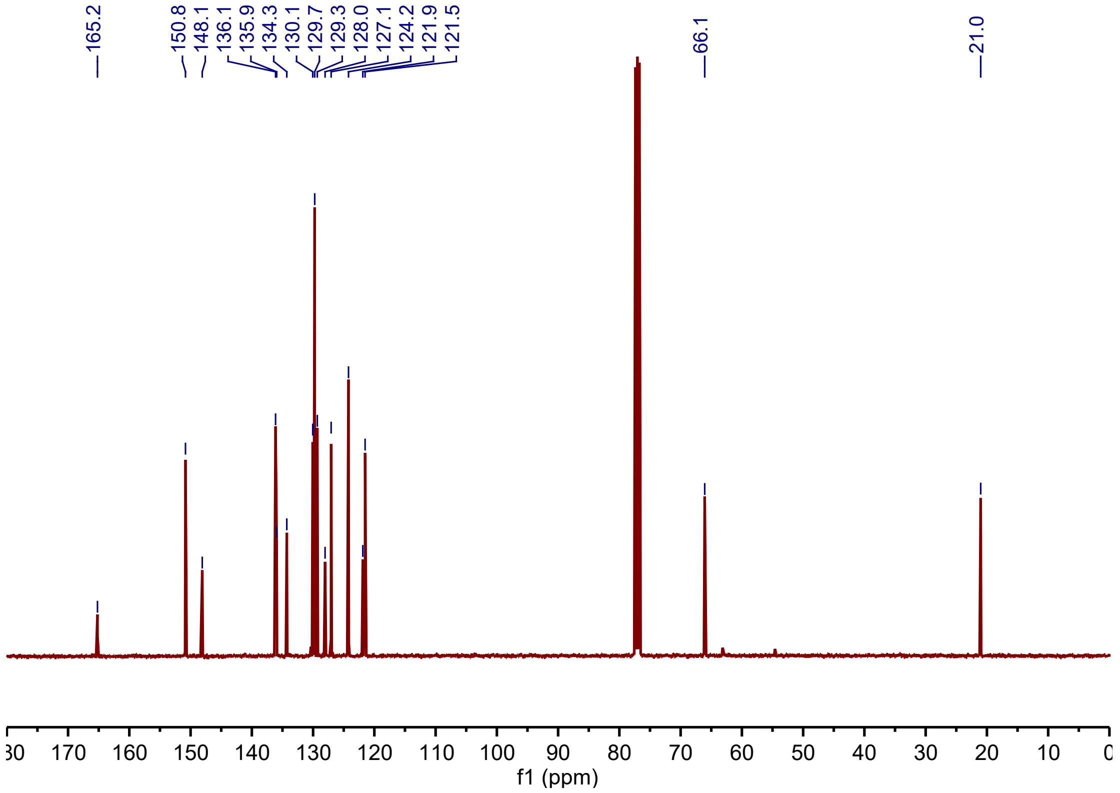

[1]H NMR spectrum(top) and [13]C NMR spectrum(bottom) of **DQmB**.
